# The cellular representation of temperature across the somatosensory thalamus

**DOI:** 10.1101/2024.02.13.580167

**Authors:** Tobias Leva, Clarissa J. Whitmire, Ilaria Sauve, Phillip Bokiniec, Charlene Memler, Bibiana M. Horn, Mikkel Vestergaard, Mario Carta, James F.A. Poulet

**Author notes:** These authors contributed equally (Tobias Leva, Clarissa J. Whitmire).

## Abstract

Although distinct thalamic nuclei encode sensory information for almost all sensory modalities, the existence of a thalamic representation of temperature with a role in thermal perception remains unclear. To address this, we performed high-density electrophysiological recordings across the entire forelimb somatosensory thalamus in awake mice, and identified an anterior and a posterior representation of temperature that spans three thalamic nuclei. We found that these parallel representations show fundamental differences in the cellular encoding of temperature which reflects their cortical output targets. While the anterior representation encodes cool only and the posterior both cool and warm; in both representations cool was more densely represented and showed shorter latency, more transient responses as compared to warm. Moreover, thalamic inactivation showed a major role in thermal perception. Our comprehensive dataset identifies the thalamus as a key structure in thermal processing and highlights a novel posterior pathway in the thalamic representation of warm and cool.

## Introduction

The thermal system is a fundamental sensory system required for survival and mammals are acutely sensitive to changes in skin temperature ^1,2^. The cortex is necessary for thermal perception ^3–5^, implying a central role for thalamus in thermal processing. However, extremely few thalamic neurons have been identified that show graded encoding of temperature, particularly warm ^6–17^. Moreover, in contrast to the cortex ^4,5^, previous attempts at thalamic inactivation have shown a weak impact on thermosensation ^18,19^ and no effect on thermoregulatory behavior ^20^. While understanding the thalamic encoding of sensory information is critical for a mechanistic understanding of perception ^21^, whether and how thermal information is processed by the thalamus was unclear.

Classical models suggest that all somatosensory information, including thermal, is forwarded to the primary somatosensory cortex (S1) via the ventroposterior lateral nucleus (VPL) and posterior complex (PO) of the thalamus ^22^. Craig et al., suggested an alternative model whereby thermosensory information is represented in the posterior ventral medial posterior nucleus (VMpo), which forwards thermal information to the posterior insular cortex (pIC) ^11,23^. The rodent homologue of primate VMpo has been proposed to be PoT ^24^ and is known to be a central target of spinothalamic neurons ^25^. PoT neurons respond to painful stimuli ^24,26,27^, but innocuous thermotactile encoding in PoT has not been addressed. Overall, because there has not been a comprehensive investigation into thalamic cellular encoding of temperature, there is no consensus about the location of the representation of temperature in the thalamus and these models remain debated ^28–31^.

Here we performed high density electrophysiological recordings in VPL, PO and PoT to generate a functional and spatial map of the thalamic cellular encoding of temperature. Using pharmacological inactivation in trained mice we show the thalamus is critically required for thermal perception. Our data highlight the importance of a novel representation in posterior regions of VPL and PO as well as PoT for thermal processing. Together we reconcile contrasting models of thermal processing in the thalamus and lay a platform for future work addressing neuronal mechanisms of thermal encoding and perception.

## Results

### Thalamic cellular tuning to thermal stimuli is defined by spatial location

Using high-density extracellular probes, we recorded from VPL, PO and PoT in awake head-restrained, paw-tethered mice (Fig. 1A, Extended Data Fig. 1). Thermal stimuli (2 s duration, 32°C +/-10°C) were delivered using fast temperature transients (75°C/s) via a Peltier element stimulator positioned on the plantar surface of the forepaw. A force-feedback tactile stimulator was positioned on the dorsal surface of the forepaw that delivered tactile stimuli (2s, 40 Hz) as well as providing a sensitive monitor of paw movement. Post-hoc reconstruction of electrode tracks, using histological and electrophysiological markers, allowed confirmation of recording sites in VPL, PO and PoT (Fig. 1B, Extended Data Fig. 2).

**Figure 1:**
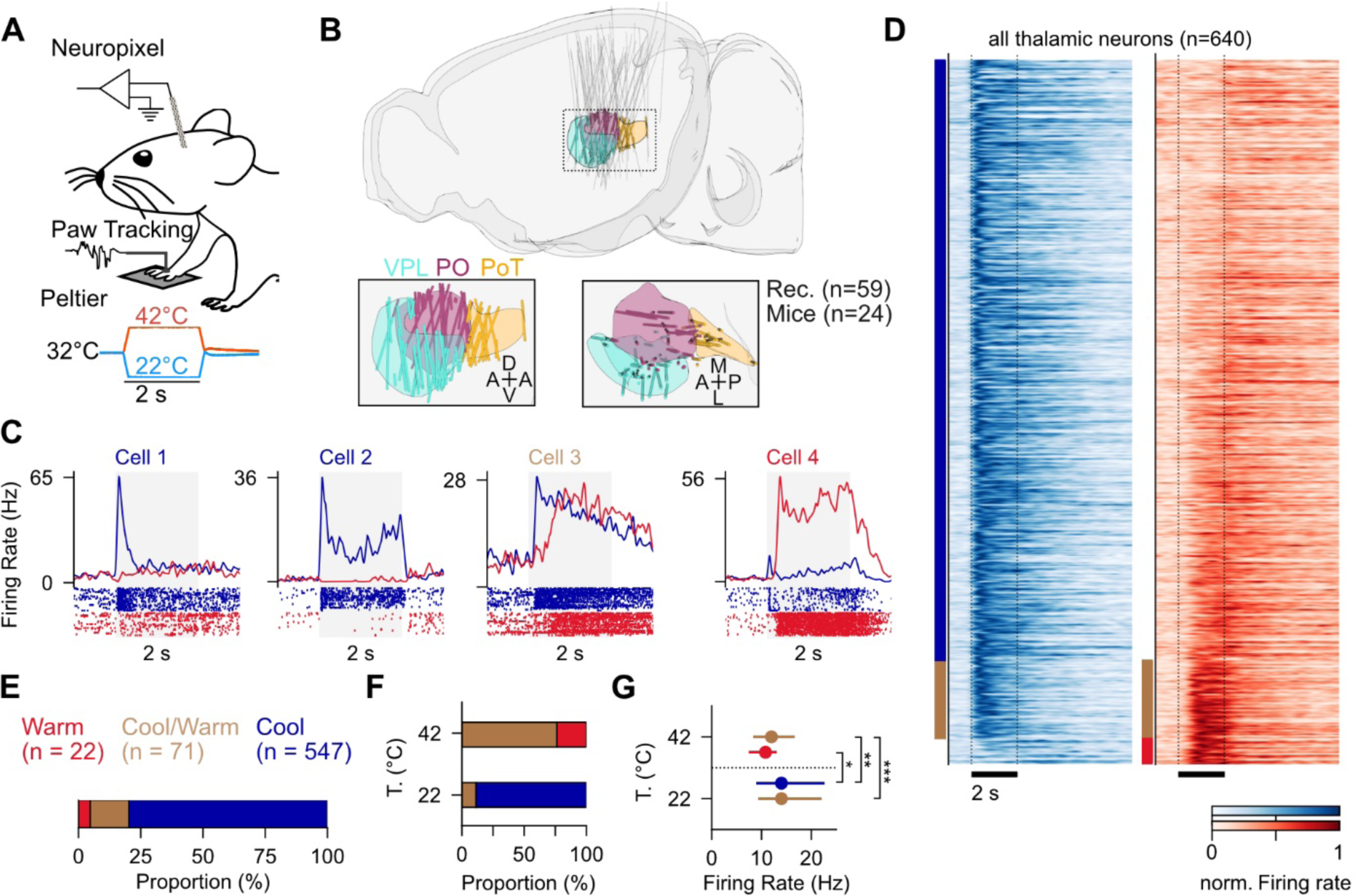
The thalamic representation of cool is over represented relative to warm. (**A**) Schematic of awake mouse during extracellular recordings from thalamus using high density multielectrode arrays (Neuropixel) during thermal stimulation, tactile stimulation, and movement tracking. (**B**) Reconstructed probe trajectories for all sessions (n = 59 probes; 24 mice). Ventral posterolateral nucleus (VPL; cyan), Posterior nucleus (PO; maroon) and Posterior triangular nucleus (PoT; yellow). Insets display magnified view of trajectories in sagittal (left) and horizonal (right) views with electrode position within the boundaries of the regions of interest indicated by a colored dot. (**C**) Example single units demonstrating the diversity of thermal responses in thalamus. Text color indicates response classification as indicated in E. (**D**) Peristimulus time histogram for each thermally responsive unit in response to a 22°C (left; blue) and 42°C (right; red) stimulus. Index bar indicates thermal responsivity for each unit as indicated in E. (**E**) Proportion of thalamic units responsive to temperature (n = 547 cool-only units, blue; 22 warm-only units, red; 71 cool-warm responsive units, tan). (**F**) Proportion of warm (top: 42°C) and cool (bottom: 22°C) responses from cool-warm units. (**G**) Peak stimulus evoked response for warm (top: 42°C) and cool (bottom: 22°C) responses from each response classification as indicated in E.

We observed cells responsive to thermal stimuli (Fig. 1C and D) across all three thalamic nuclei. Thermally responsive neurons were defined as those showing significant changes in firing rate to thermal stimuli before stimulus-evoked movement of the forepaw (Extended Data Fig. 3 A to D). The majority of temperature-sensitive neurons showed an increase in firing rate to thermal stimulation and a minority showed suppression (1% warming suppressed, 3% cooling suppressed; Extended Data Fig. 4A to E). A thermal tuning index ((max cooling – max warming) / (max cooling + max warming)) estimated the relative sensitivity of a neuron to cooling and warming and classified response types as cool only (Cool), warm only (Warm) or responsive to both cooling and warming (Cool/Warm) (Extended Data Fig. 5A). Tactile stimulation further identified tactile and thermo-tactile responsive units (Extended Data Fig. 6A to E). Notably, there was an overrepresentation of Cool units compared to Warm (Fig. 1E), and the majority of warming-responsive cells were classified as Cool/Warm (Fig. 1F). Across the population, cooling evoked higher firing rates than warming independent of response type (Fig. 1G). Together, these data show that both warming and cooling drive thalamic cellular responses and highlight a stronger thalamic representation of cooling than warming.

The thermally sensitive cortical regions display functionally distinct representations of temperature. Both warming and cooling are represented in pIC, but only cooling in S1 (*4*–*6*, *25*). Therefore, we hypothesized that thalamic nuclei will show thermal tuning specificity with respect to their cortical target. Retrograde tracing from the thermal cortices (Fig. 2A) shows separability of thalamic neurons by cortical projection target along the anterior-posterior axis (*23*). Posterior regions of VPL and PO as well as PoT contain cells projecting to pIC, whereas more anterior regions of VPL and PO project to S1 (Fig. 2B, 2C). PoT does not project to S1 (Fig. 2D). Therefore, we mapped the anatomical location of each thermally responsive thalamic unit onto their thalamic location using the reconstructed electrode contact position (Fig. 2E to J). To better link spatial position to cortical projection target, we defined an anatomical boundary (Fig. 2B, C, F, G, I, J, vertical dashed lined) between anterior and posterior to delineate the thalamic nuclei based on the anatomical data (anterior VPL anterior PO and PoT to pIC and posterior VPL and PO to S1, Fig. 2D).

**Figure 2:**
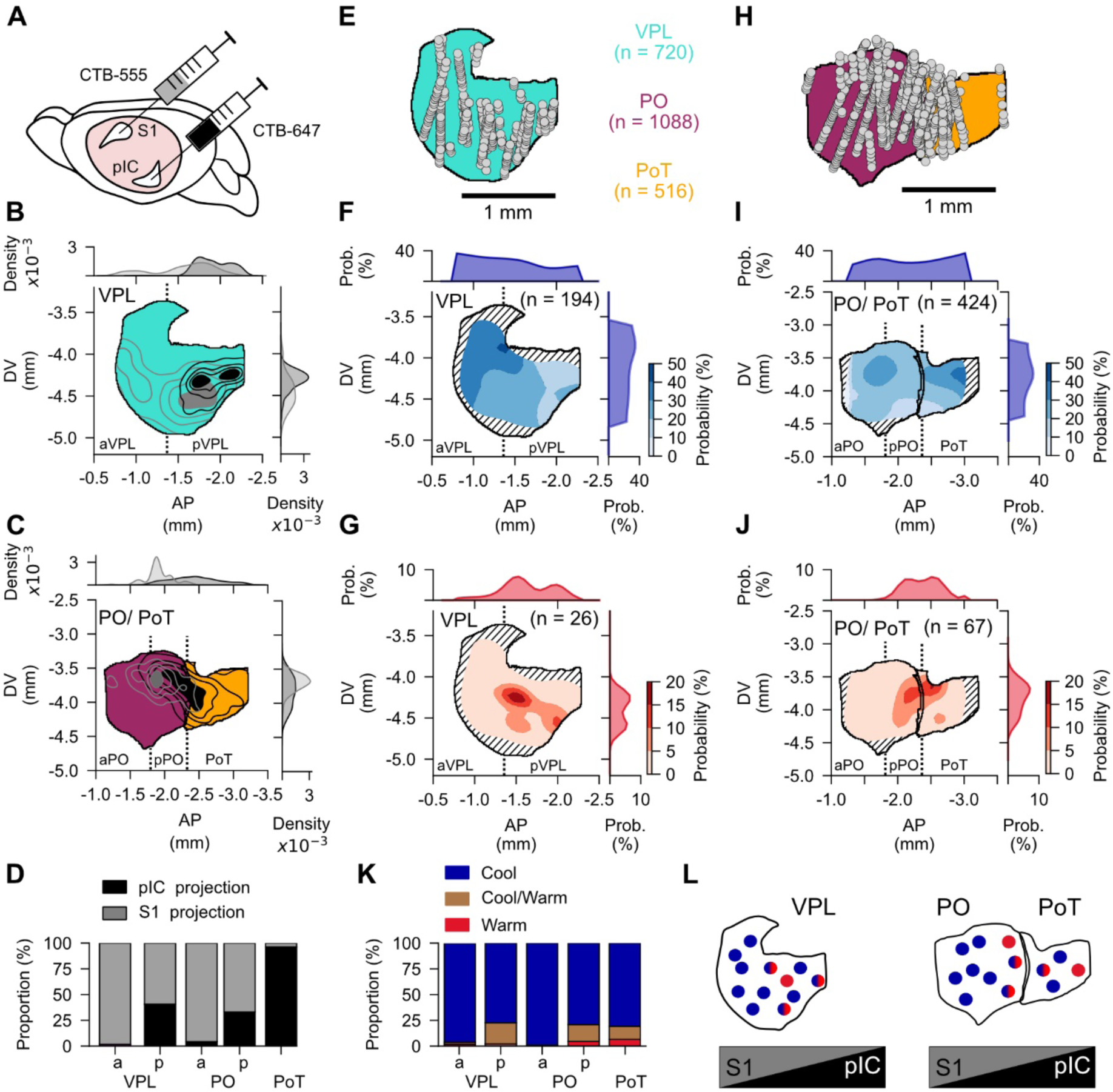
Thalamic representation of cool and warm varies by spatial location. (**A**) Schematic of retrograde anatomical tracing from thermosensory cortical regions (data reanalyzed from^32^). (**B**) Reconstructed position of retrogradely labeled thalamocortical neurons located within the VPL projecting to S1 (grey) or pIC (black). (**C**) Reconstructed position of retrogradely labeled thalamocortical neurons located within the PO/PoT nuclei projecting to S1 (grey) or pIC (black). (**D**) Proportion of retrogradely labeled thalamocortical neurons projecting to S1 (grey) or pIC (black) relative in anterior (a) or posterior (p) regions of VPL and PO as well as PoT. (**E**) Reconstructed position of each recorded thalamic unit (grey dot) in VPL (teal) shown in sagittal projection. (**F/G**) Spatial density maps for VPL units responsive to cool (I: blue, n = 194 cool units recorded in n = 13 mice) or warm (J: red, n = 26 warm units recorded in n = 9 mice). Contour lines drawn at 10^th^, 20^th^, 30^th^, 40^th^ percentiles. Hatched infill indicates insufficient sampling to estimate density. (**H**) Reconstructed position of each recorded thalamic unit (grey dot) in PO (maroon), and PoT (yellow) shown in sagittal projection. (**I/J**) Spatial density maps for PO/PoT units responsive to cool (I: blue, n = 424 cool units recorded in n = 16 mice) or warm (J: red, n = 67 warm units recorded in n = 9 mice). Contour lines drawn 10^th^, 20^th^, 30^th^, 40^th^ percentiles. Hatched infill indicates insufficient sampling to estimate density. (**K**) Proportion of functional response-type by thalamic sub-region. (**L**) Schematic depicting spatial gradient of thalamic thermal cellular tuning and cortical projection targets.

Reconstruction of spatial probability distributions for cooling responsive neurons in VPL showed a broad distribution, with a larger proportion of neurons located in anterior VPL (aVPL, Fig. 2F). In contrast, warming responsive neurons were spatially localized in a restricted zone of posterior VPL (pVPL, Fig. 2G). Likewise, in PO and PoT, cooling responsive units were found across the anterior-posterior axis, with the highest proportion in PoT (Fig. 2I); whereas warming responses were localized at the junction between posterior PO (pPO) and PoT (Fig. 2J). The unique arrangement of warming responsive neurons was in contrast to the spatial representation of tactile and paw movement responsive units. The numbers of units responsive to tactile stimulation have a relatively even distribution across VPL, PO and PoT (Extended Data Fig. 6F to H), whereas movement units have a higher concentration in aVPL and PO compared to pVPL, pPO or PoT (Extended Data Fig. 3H and I). The functional arrangement of thermally responsive units closely aligns with the tuning of the cortical project target. The posterior regions of VPL, PO and PoT project to the warm and cool responsive pIC, whereas anterior VPL and PO project to S1 which is only responsive to cool (Fig. 2D), indicating a close match between thalamic tuning and cortical target (Fig. 2K, Extended Data Fig. 5B to D). Together these data support the hypothesis that there are two major representations of temperature across the somatosensory thalamus: an anterior representation that is cooling-selective and projects to S1, and a posterior representation that is warming and cooling sensitive and projects to pIC (Fig. 2L).

### Thalamic encoding of thermal stimulus amplitude follows an intensity model

Different models have been proposed to describe the encoding of cooling and warming in thermal primary sensory afferent neurons. The ‘specificity model’ suggests that thermally sensitive neurons are tuned to specific temperature amplitudes, whereas an ‘intensity model’ proposes that response amplitude changes monotonically with stimulus intensity (Fig. 3A) ^33^. To investigate whether these models exist in the thalamus, we delivered a range of temperatures values from a baseline, adapted temperature of 32°C and classified the units as either specificity or graded encoders (Fig. 3B and C, see methods). Overall, 98% of thermosensitive thalamic neurons follow a graded coding scheme in response to cooling stimulation (Fig. 3B and cells #1-5 in 3C), and only 2% showed specific tuning to intermediate temperature values (Fig. 3C cell # 6, Extended Data Fig. 4F to H). Notably, all warming sensitive neurons followed an intensity based encoding scheme (Fig. 3B). Therefore, intensity coding is the dominant thalamic encoding method within the innocuous thermosensory range.

**Figure 3:**
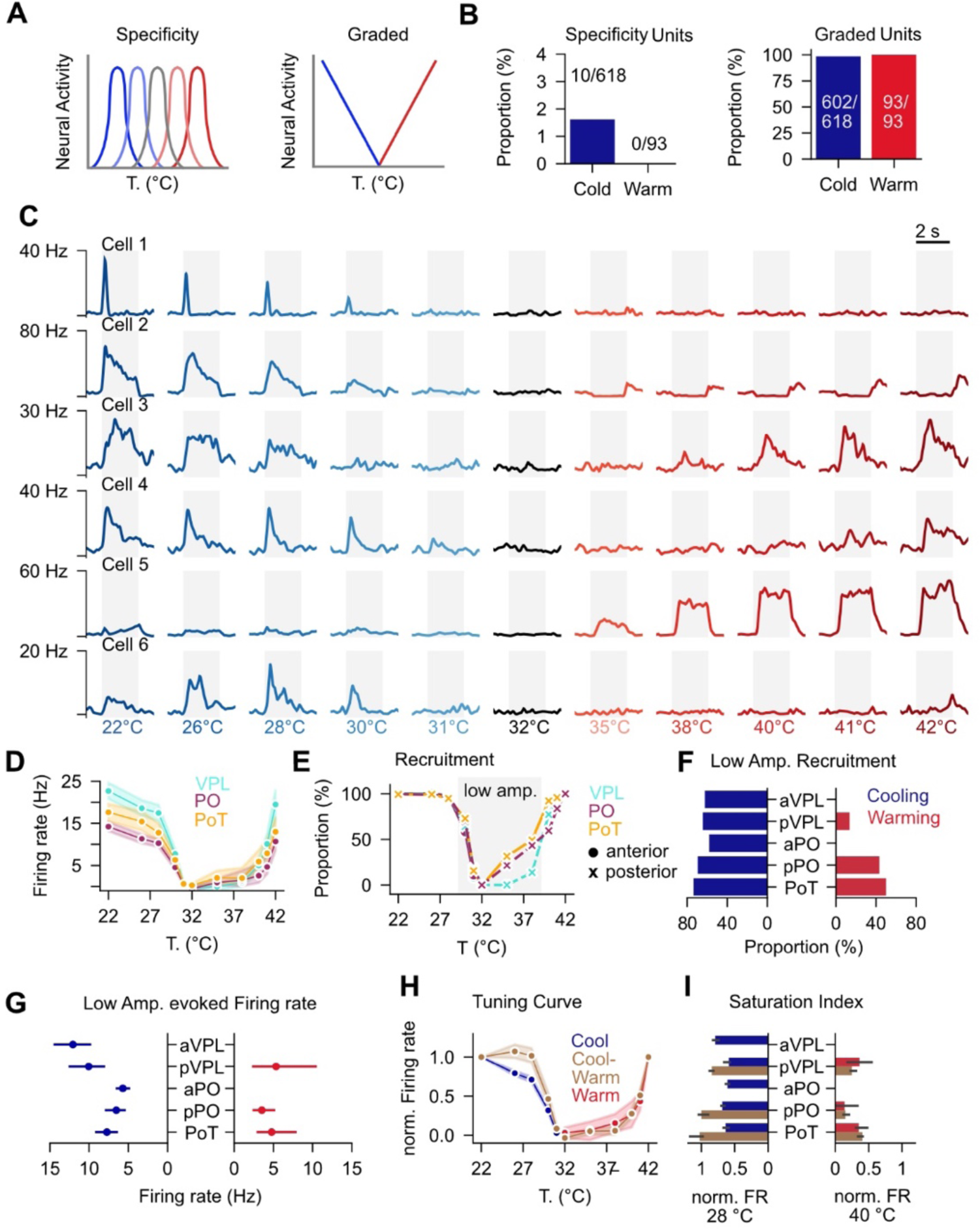
Graded cellular encoding of temperature amplitude in the thalamus. (**A**) Schematic showing thermal specificity tuning (left) and graded thermal (right) coding schemes. (**B**) Proportion of neurons that follow a specificity model (left) or graded model (right) in response to cooling (blue) or warming (red). (**C**) Example units PSTH for graded encoding of temperature. Each row is one example unit and each column is a single stimulus intensity. Grey shading indicated stimulus duration. (**D**) Stimulus evoked response amplitude for each stimulus intensity by thalamic nuclei. Cooling responsive neurons are shown for cooling stimuli (VPL: n = 194 neurons, PO: n = 283 neurons, PoT: n = 141 neurons) and warming responsive neurons are shown for warming stimuli (VPL: n = 28 neurons, PO: n = 38 neurons, PoT: n = 29 neurons). (**E**) Proportion of neurons recruited at 20% of the maximum stimulus evoked firing rate by thalamic nucleus. Warming and cooling responses were normalized independently. (**F**) Proportion of neurons recruited at low amplitude cooling (left) or warming (right) by sub-nucleus. (**G**) Low amplitude cooling (left) or warming (right) stimulus evoked firing rate by sub-nucleus. (**H**) Normalized tuning curves by response type classification. Warm and cool responses were normalized independently. (**I**) Saturation Index for cool, cool-warm, and warm response types by thalamic sub-nucleus.

While both warming and cooling can be described by an intensity model, our data showed a clear difference in the shape of the cooling and warming thermal tuning curves in all nuclei (Fig. 3D). Cooling response curves show a sharper rise and prominent plateau compared to a more graded profile for warming across nuclei and within subregions (Fig. 3D, Extended Data Fig. 7A). To quantify the sensitivity of thalamic neurons to low amplitude warming and cooling, we computed the proportion of neurons recruited at 20% of the maximum stimulus evoked firing rate at each stimulus amplitude (Fig. 3E). This analysis showed that cooling recruited more neurons at low amplitudes than warming across all nuclei independent of functional cell response type (Fig. 3F, Extended Data Fig. 7B). Moreover, firing rates evoked by low amplitude stimuli were lower for warming than cooling, with VPL showing the highest cooling evoked firing rates (Fig. 3G). Taken together, this suggests that the thalamic representation of temperature is more sensitive to cooling than warming, consistent with higher perceptual acuity for cooling relative to warming in both humans and mice ^1,2^.

Increased sensitivity to sensory stimuli can also lead to an earlier saturation of the sensory response. Saturation of a sensory neuron limits the dynamic range and therefore the capability to faithfully encode high amplitude stimuli. To quantify the saturation of thalamic neurons, we normalized the tuning curves to the response amplitude at maximum cooling and warming respectively (Fig. 3H) and defined the saturation index as the normalized firing rate at an intermediate stimulus intensity (28°C for cooling and 40°C for warming, Fig. 3I). Cooling responses were more saturated than warming responses regardless of the functional cell type classification (Cool, Warm, Cool/Warm) or thalamic subregions, indicating that cooling responses have a smaller dynamic range (Fig. 3H and I). Unexpectedly, we found that the cooling saturation was dependent on the functional cell type classification. Cool/Warm neurons, which exist solely in the posterior thalamic representation, saturate at lower cooling amplitudes compared to Cool only neurons (Fig. 3H and I). A lower saturation point can lead to greater detectability of thermal stimuli at the expense of discriminability. This indicates that Cool/Warm neurons detect thermal identity while Cool-only neurons are responsible for discrimination of cooling intensity.

### Temporal encoding of temperature varies with thermal tuning and spatial location

Recordings of primary sensory afferents and cortical neurons have revealed differences in thermal evoked temporal dynamics with warming eliciting longer latency, sustained responses while cooling elicits shorter latency responses that can be either sustained or transient ^1,4,5,34–38^. Consistent with these findings, population analysis of thalamic responses shows a shorter response latency with an earlier peak time to cooling than warming (Fig. 4, A and B; Extended Data Fig. 8A to D). Interestingly, these properties are consistent within cells that respond to both cool and warm (Extended Data Fig. 8E and F). This suggests that differences between cool and warm response dynamics are due to a merging of hard-wired afferent inputs within thalamic neurons, rather than the result of thalamic processing.

**Figure 4:**
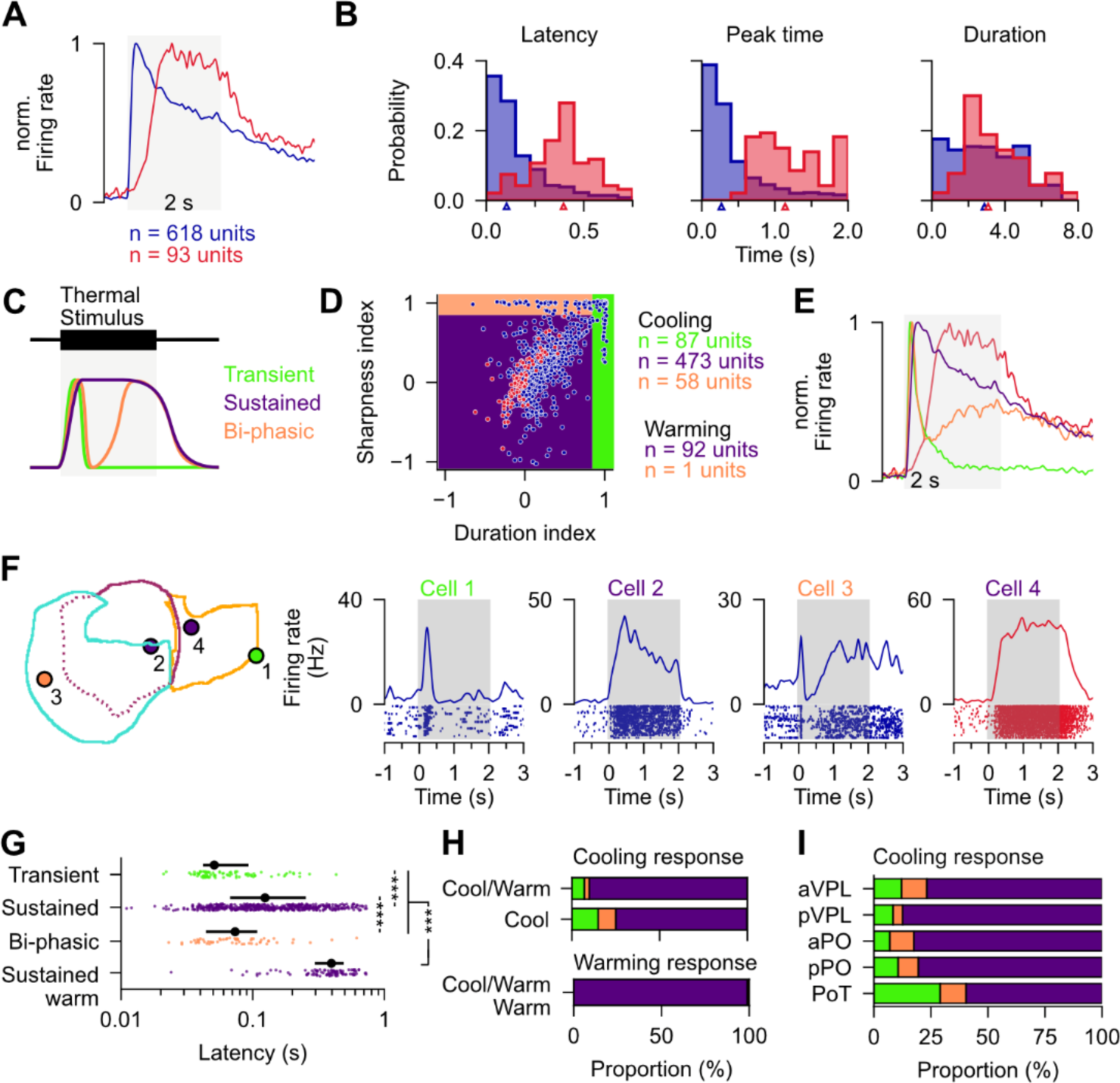
Temporally sharpening of thermal encoding along the anterior-posterior axis. (**A**) Normalized stimulus-evoked PSTH to 22°C (blue) and 42°C (red). Stimulus duration indicated by grey shading. (**B**) Probability distributions for stimulus response onset (left), peak time (middle), and duration index (right) in response to cool (22°C, blue, n = 618 neurons) and warm (42°C, red, n = 93 neurons). (**C**) Schematized response dynamic subtypes for cool (transient: green, biphasic: orange, sustained: purple) and warm (sustained: purple). (**D**) Scatterplot of the cooling (blue) and warming (red) evoked response dynamics parameterized as the duration index and the sharpness index. Transient responses have a duration index > 0.85; Biphasic response types have a sharpness index > 0.85 (and duration index < 0.85). Sustained responses have a duration & sharpness index < 0.85. (**E**) Normalized cooling stimulus-evoked PSTH for each response dynamic type. (transient: green, biphasic: orange, sustained: purple). Stimulus duration indicated by grey shading. (**F**) Left, spatially reconstructed sagittal locations and, right, PSTH with corresponding spiking rasters of three example cool responsive neurons (Cells 1-3) and one example warm responsive neuron (Cell 4). (**G**) Latency as a function of response dynamic type. Colored dots denote individual units and black dots denote the population distribution (median +-IQR). (**H**) Proportion of units classified as transient (green), biphasic (orange), or sustained (purple) response dynamics by thermal response type. (**I**) Proportion of cooling responsive units classified as displaying transient (green), biphasic (orange), or sustained (purple) response dynamics by thalamic nucleus.

Thermosensory response dynamics of afferent neurons have been further defined into categories of transient, sustained, and bi-phasic responses ^39^ (Fig. 4C). To classify whether these subtypes exist in the thalamus, we defined functional cell types using indices for response duration and sharpness. A duration index value of 1 indicates a cell with a sensory response contained within the first half of the sensory stimulus while a duration value of -1 indicates a cell with a sensory response contained only with the second half of the sensory stimulus. A sharpness index value of 1 indicates a cell with a peak in the first 500 milliseconds of the stimulus response. Cells with high duration indices were defined as transient. Cells that had high sharpness index and low duration index were defined as bi-phasic (Figure 4D, Extended Data Fig. 8G-H), as confirmed by the population PSTH by dynamic subtype (Fig. 4E). Within the warming responsive population, nearly every neuron (n = 92 / 93 neurons) was classified as sustained, with only one neuron classified as bi-phasic. In contrast, cooling responsive neurons contained all three categories including transient (14%), bi-phasic (9%) and sustained (77%) response dynamics (Fig. 4F). We observed that transient and biphasic cooling responsive neurons had the shortest response latency, whereas sustained cells response to cooling has a broader distribution of latencies but is still significantly faster than the sustained warming response (Fig. 4G). Across functional response types, the Cool only population contained more transient and bi-phasic responses than Cool/Warm (Fig. 4H). Spatially, we observed the lowest proportion of sustained cells and the highest proportion of transient response type in PoT (Fig. 4I). Further subdivision of cells within the sustained population indicate that the highest proportion of short duration and short latency sustained responses are also found in PoT, further emphasizing the differences in response dynamics across thalamic subnuclei (Extended Data Fig. 8I to N). A high proportion of transient response types were also observed in PoT and posterior PO to tactile stimulation of the forepaw (Extended Data Fig. 9), suggesting that fast, transient temporal response profiles in the posterior representation are nucleus-rather than modality-specific.

### The thalamic representation of temperature is required for thermal perception

Early human lesion studies indicated a role for the thalamus in thermal perception ^40^, but experimental manipulations have only shown mild deficits in thermal perception ^18,19,41,42^. To address this, we performed pharmacological inactivation of the thalamus in mice performing a thermal perception task ^5^. Mice were trained on a Go/NoGo task to report either 10°C warming or 5°C cooling delivered from an adapted temperature of 32°C (Fig. 5 A and B). Once mice were trained (Hit rate > 70%, FA-rate < 30%), we injected ∼100nl of the GABA-agonist muscimol into the thalamus. Following testing, fluorescently labeled pipette tracts were reconstructed anatomically and the tip location was used to define the center of the injection site (Extended Data Fig. 10A, 10C).

**Figure 5:**
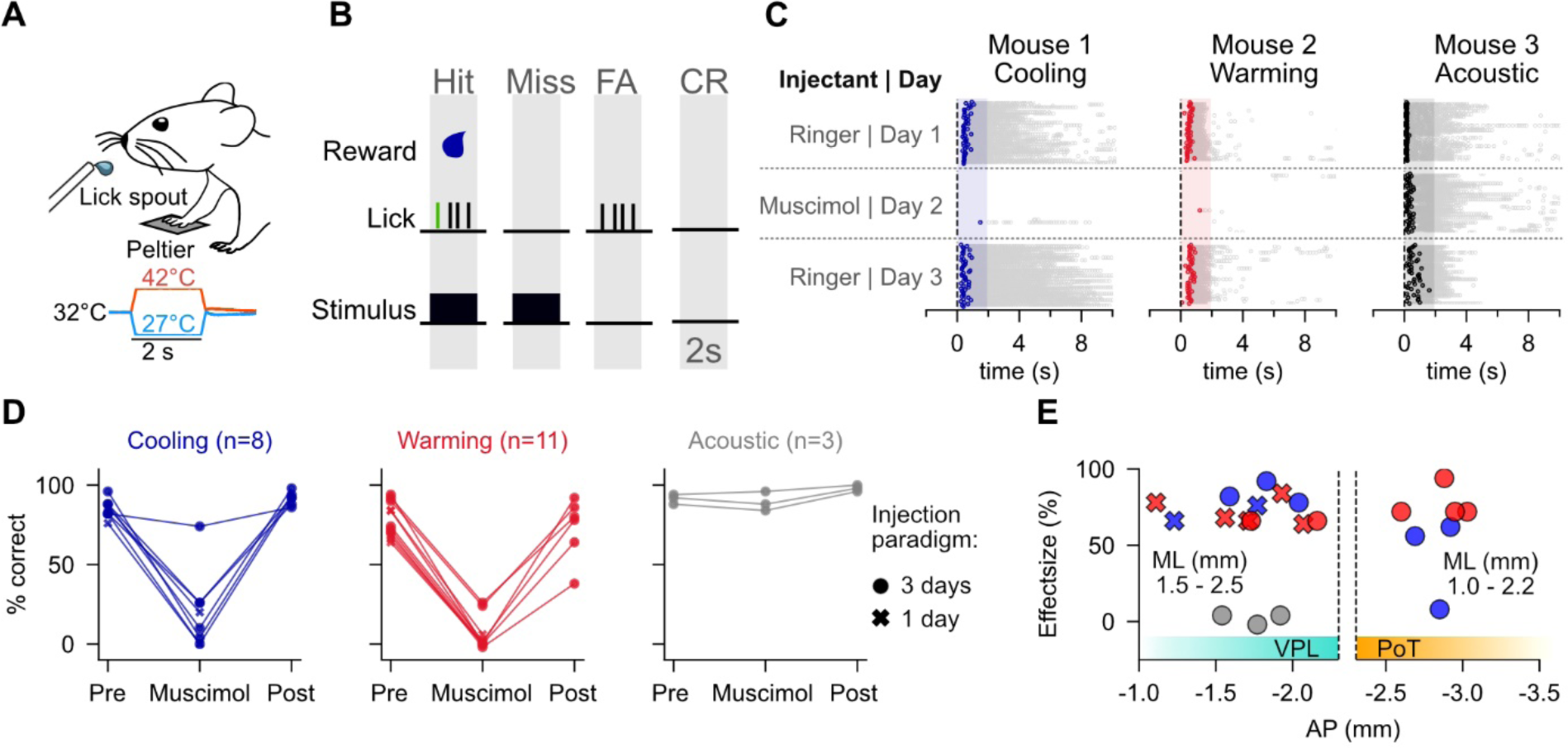
Thermal perception requires the thalamus. (**A**) Left, cartoon showing go/no-go thermal or auditory detection task; (**B**), behavioral response categories. (**C**) Experimental timeline (left) describes injectant for control (Day 1), manipulation (Day 2), and recovery (Day 3). Example data for three mice trained on the three-day experimental timeline. Lick rasters shown for mice trained on a cooling (Mouse 1; blue, left), warming (Mouse 2; red, center), and auditory (Mouse 3; black, right) task. (D). Behavioral performance of all mice in response to muscimol injection during cooling (n = 8 mice), warming (n = 11 mice), or auditory (n = 3 mice) tasks shown as the percentage correct. Filled circle marker indicates that the mouse underwent a three-day injection paradigm as outlined in panel (C), while a cross mark indicates that the mouse underwent a single-day injection of muscimol without a recovery session, see Methods. (**E**) Effect size (change in hit rate on Day 1 vs Day 2) for cool (blue), warm (red) and acoustic (grey) task performance following injections in posterior regions of VPL (left) or PoT (right). Marker indicates whether the mouse underwent a three-day injection paradigm as outlined in panel C (filled circle) or a single-day injection paradigm of only muscimol (x marker) without a recovery session. AP, Anterior-posterior; ML, mediolateral.

Injection of muscimol into somatosensory thalamus suppressed licking responses to the warming or cooling stimuli but did not alter performance on an acoustic detection task (Fig. 5 C and D, Extended Data Fig. 10B). These data therefore indicate that somatosensory thalamic inactivation selectively altered thermal perception rather than the ability to perceive any sensory stimulus or licking movements. Across mice, muscimol injection had a marked effect on behavioral performance (Fig. 5D), primarily attributed to a reduction in hit responses (Extended Data Fig. 10B, 10D). The difference in behavioral performance on the muscimol injection day relative to the previous day, defined as the ‘effect size’, was significant across the anterior-posterior somatosensory axis (Fig. 5E, Extended Data Fig. 10E, 10F). Together, these data suggest that both the anterior and the posterior thalamic representation of temperature play a central role in thermal perception.

## Discussion

The thalamus is required for perception of almost all modalities of sensory information, however little is known about the thalamic representation of temperature. To address this, we performed a comprehensive mapping of the cellular encoding of temperature across the somatosensory thalamus. We show that cool and touch responsive units are widely distributed across the thalamus, whereas warm responses are located only in posterior regions of VPL and PO as well as in PoT. Moreover, our inactivation experiments in mice performing a thermal perception task show that the thalamus is required for both warm and cool perception. Our data help resolve competing models of the thalamic representation of temperature and suggest the presence of two parallel somatosensory thalamocortical pathways for temperature encoding: an anterior pathway encoding cool, and a posterior pathway encoding cool and warm.

Previous investigations into the cellular encoding of innocuous temperature in the thalamus have almost exclusively investigated cool and tactile processing in anterior regions of the somatosensory thalamus ^7,9,10,13,17^. Our data show an almost complete absence of warming responses in anterior VPL. This aligns well with the absence of identified warm sensitive units in limb regions of the human thalamus ^16^. Furthermore, it matches the cool selective tuning of S1, which is the cortical target of anterior regions of VPL and PO. We show that warm responsive units are present in the thalamus, but only in posterior regions of VPL and PO as well as in PoT. This supports the proposal of a posterior thalamic representation for temperature encoding in primates ^11^. It also matches the tuning of the cortical target of these posterior regions, pIC, which is sensitive to both warm and cool.

Alongside a representation of warming, the posterior representation in PoT had fewer movement responsive units, higher levels of unit recruitment at low amplitude thermal stimuli and proportionally more transient response types than anterior regions. Little is known about PoT, but it has been traditionally considered to play a role in pain and itch sensation rather than innocuous thermal processing ^24,27,43–45^. PoT is heavily innervated by spinothalamic tract neurons ^25^, that are sensitive to temperature and noxious stimuli ^46,47^, and projects to the primary thermal cortical representation ^24,32^ in the pIC which is both warm and cool sensitive. Together with its responsiveness to warm and robust impact on thermal perception, our study highlights PoT as a central node in the thermal pathway and support the hypothesis that it is the rodent homologue of VMpo ^24^.

In 1882, Blix identified spatially discrete spots in the skin that could evoke warm or cool sensations suggested a functionally segregation of warm and cool processing ^48^. In contrast, recordings of thermally sensitive cortical neurons have demonstrated individual neurons responding to both warm and cool. We also observe cool-warm neurons in the thalamus. Intriguingly, as in the cortex, the encoding features of warm and cool were markedly different even within individual thalamic neurons sensitive to both cool and warm. Warm responses were longer latency and more sustained than cool, which was represented by both sustained and transient responses. One possibility is that longer latency, sustained responses are driven by thermally sensitive c-fibers, whereas shorter latency, transient thalamic responses to cool could be driven by the thinly myelinated a-delta fibers ^1,34,49–51^. The tuning curves for warm and cool also were also distinct (Fig. 3D). Cool showed a more rapid recruitment and higher response rates to low amplitude stimulation than warm, closely resembling tuning curves from cortical neurons in pIC ^4^. While most thermally sensitive neurons showed excitatory responses to thermal stimuli, we also observed a smaller population of suppressed responses that resembled classical afferent thermoreceptors (Extended Data Fig. 4) ^52^. Taken together these data suggest some thermal afferent information is directly copied from the periphery via the thalamus to the cortex. It moreover indicates that warm and cool are not just two ends of a single modality spectrum but appear more like different sub-modalities of temperature ^4^.

There were also some notable differences of thalamic thermal responses to afferent input or cortical neurons. For example, our thalamic data match afferent data that show a sparse encoding of warm compared to cool ^8^, yet cortical layer 2/3 neurons show a more similar proportion of warm and cool responsive cells ^4^. Further work could test whether this results from differences in recording techniques, an additional source of warm input to the cortex or an integration of thalamic inputs. Moreover, in contrast to a report that investigated mouse hindpaw afferent neuron thermal encoding ^33^, we only observed a tiny number of specificity units in the thalamus with preferred temperature responses. In recent work we reported ‘thermometer’ like cells in the cortex that encode stimulus amplitude independent of stimulus direction and were characterized by low threshold responses to warming and selective responses for low amplitude cooling ^4^. We did not observe this response type in the thalamus and, as these responses have also not been reported in the periphery, it raises the possibility of a different input pathway to the cortex, or the generation of thermometer cell responses within the cortex.

While touch and cool were present across all thalamic regions tested, warm is absent from anterior regions. One possibility for the overrepresentation of cooling in the thalamus could be due to the natural thermal statistics of the somatosensory environment. In homeothermic mammals, surfaces are inherently cool-biased and the more prevalent cool responses could be expect given the higher likely hood of handling cool than warm objects. It is also possible that additional cool information provides information about surface texture during haptic exploration. For example, in mammals a sensation of wetness is thought to be an integration of cool and tactile information rather than the result of a specific wetness afferent input ^50^. The presence of warm alongside cool in the posterior regions of VPL and PO as well as PoT, alongside the presence of warm and cool in the pIC, suggest that the posterior pathway is critical for temperature representation.

Our findings bring together disparate findings about anterior and posterior representations of temperature in the somatosensory regions of the thalamus. This study could guide future investigations into functional differentiation of the thermosensory pathways and their involvement in somatosensation more broadly.

### Online methods

All experiments were approved by the Berlin Landesamt für Gesundheit und Soziales (LAGeSo), and carried out in accordance with European animal welfare law. Adult male and female C57BL/6J mice were housed under 12-hour light/dark cycles and provided with ad libitum food and water.

### Surgery

Mice were anesthetized with an intraperitoneal injection of Ketamine (120 mg/kg) and Xylazine (10 mg/kg). Body temperature was maintained at 37°C throughout the surgical procedure using a heating pad (FHC). Mice were fixed in a stereotaxic frame (Narishige) and the skull was exposed. The stereotactic coordinates for thalamic extracellular recordings were marked on the skull. An aluminum bar was glued to the parietal bone of the right hemisphere (Loctite 401, Henkel) and a recording chamber was made around the recording sites (C&B, Metabond). The exposed skull was covered with a silicone elastomer (Kwik-cast) to protect the surface until the recording session. After the surgery, mice were placed on a heating blanket until they recovered from anesthesia. Drinking water in the home cage was supplemented with Metamizol (200 mg/kg, Ratiopharm) for 2 – 3 days for pain management.

### Extracellular Electrophysiology

Following recovery from surgery, mice were habituated to head fixation and paw fixation across multiple sessions. Mice were exposed to the full thermotactile stimulus set on 1-2 sessions prior to the recording session to habituate mice to the stimulation paradigm. On the day of recording, the mice were lightly anesthetized with isoflurane and a small (0.5 mm) craniotomy was performed over the thalamic target recording regions. Mice were allowed to recover from this procedure for 1-2 hours before the recording session began.

Mice were head-fixed and paw-tethered during recordings. The recording chamber was filled with Ringer’s solution. A silver wire coated with chloride served as the reference electrode. Neural recordings were performed with Neuropixels probes (Neuropixels 1.0, IMEC) with 383 active channels. The channels closest to the tip were selected for recordings, which span 3.84 mm of tissue. Neuropixels probes were coated with fluorescent dye (DiI, DiI-CM or DiD; 1mM in 100% Ethanol) to localize the probe position during the ex-vivo imaging of fixed brain-slices. Electrophysiology data was acquired at 30 kHz sampling rate and 500x gain for the spike band and 2.5 kHz sampling rate and 250x gain for the LFP band (Hardware: National Instruments, Software: Open Ephys GUI, https://open-ephys.org/gui).

One or two probes were mounted on a 3-axis Micomanipulator (NewScale, https://www.newscaletech.com/multi-probe-micromanipulator/). Target coordinates (A/P, M/L, D/V) were translated into absolute micromanipulator coordinates (NewScale MPM-VCS system). Each probe was manually lowered until the probe tip was in contact with the pia surface. The position of each manipulator was saved and the probes were lowered until the first spikes appeared on the electrodes closest to the tip. After visual confirmation of spiking, the probes were automatically lowered with a rate of 200 µm/min until a depth of 2.5 mm was reached. The insertion rate was reduced to 100 µm/min until the target position was reached. Probes were allowed to settle for five minutes until the recording started.

During the recording session, the awake, head-restrained mice were exposed to thermal, vibrotactile, and acoustic stimuli. The right forepaw was tethered to a custom thermal stimulator (QST.Lab, T08, https://www.qst-lab.eu/probes) to ensure consistent thermal stimulation throughout the experiment. Vibrotactile stimuli (Aurora Scientific) were delivered to the back of the tethered forepaw. Paw movement was recorded using the same device as for vibrotactile stimulation. Paw movements were recorded at a 1 kHz sampling rate. All data was synchronized to sensory stimulation using custom software (Python) to trigger stimulus onset. The sensory stimulation paradigm was delivered through National Instruments hardware using custom software (Python).

At the end of the recording session, mice were transcardially perfused with paraformaldehyde (4% PFA), and the brains were extracted. Brains were postfixed overnight in PFA before washing in a phosphate-buffered solution. Brains were sliced (100 μm sections) and mounted in DAPI media. Brain slices were imaged on an upright fluorescent microscope (Zeiss, Axio Imager A.2). Probe tracks were reconstructed anatomically ^53^. The position of the deepest contact along the probe track was corrected based on established anatomical markers including spike waveform classification in inhibitory nuclei, spontaneous firing rates, and known lack of cell bodies in white matter tracts and ventricles. Electrodes were classified as belonging to one of the thalamic nuclei of interest if it was located within 100 μm of the region border.

### Behavioral experiments

Water-restricted, head-fixed mice were trained to perform a go/no-go stimulus-detection task using thermal or acoustic stimuli. Mice were presented with a randomly timed sensory stimulus (thermal or acoustic) and received a water reward (2-4 µl) upon at least one lick of a water spout within a 2 sec window of opportunity from stimulus onset. No negative reinforcement was provided for incorrect behavior and no reward was given for a correct rejection. Control of stimulus and reward timings and data collection were performed using custom-written software in Labview (National Instruments, USA)

#### Thermal perception task

The behavioral paradigm consisted of an acclimatization, pairing, training, and a testing phase. During acclimatization, mice were given free access to water rewards from the water spout. During a pairing phase of at least 2 sessions, mice were presented with a 2 second 10 °C warming or 5 °C cooling stimulus followed by a water reward 250 ms after stimulus onset to create a positive association between stimulus and reward intake. In the subsequent training phase, mice were only given a water reward if they licked at least once within a window of opportunity between 0 – 2 s following stimulus onset. The time duration between stimulus presentations was randomized between 15-20 s. To estimate the spontaneous licking frequency of the animals, trials in which a stimulus was presented (stimulus trials) were interleaved with catch trials, in which no stimulus was delivered. The number of stimulus trials and catch trials were equal in every experiment and were pseudo-randomized. A licking response during a stimulus presentation trial was counted as a ‘hit’. Failure to lick during the stimulus presentation the trial was counted as a ‘miss’. If the animal licked within the window-of-opportunity in a ‘catch trial’, this trial was counted as a ‘false-alarm’. To assess the animal’s performance, the hit rate and false-alarm rate were calculated. The hit rate was defined as the number of hits divided by the total number of stimulus trials, and the false-alarm rate was defined as the number of false alarms divided by the total number of catch trials. If the hit rate was > 70% and false-alarm rate was < 30% on two consecutive days, the animal entered the testing phase.

Testing included the same protocol as in the training phase above, but included thalamic manipulations with Ringer’s solution injection as a control or Muscimol injections to inactivate the thalamus. Injections were performed in the VPL or PoT using stereotactic coordinates. Mice underwent either a three-day injection paradigm (Day 1: Ringer’s, Day 2: Muscimol, Day 3: Ringer’s) or a single-day injection paradigm (Muscimol). On the first day of testing, a single craniotomy (∼500 µm diameter) was performed under isoflurane anesthesia to access the target region with an injection pipette. Mice were allowed to recover for at least 2 hours in their home cage following the craniotomy. Then, awake mice were head-fixed and a glass pipette was lowered to a depth of 3.5-4.0 mm relative to the pia surface. In three-day injection paradigm mice, 100 nL of Ringer’s solution was ejected with a flow rate of approximately 100 nl/min. The pipette was left in place for 5 minutes and then slowly retracted. On the second and third day of the testing phase, the mouse underwent head-fixation and microinjections as described for the first day (Day 2: 100 nl Muscimol (5 mM), Day 3: 100 nl Ringer’s solution with CTB (1:1 v/v). The injection pipette was coated in a fluorescent stain (Day 2: DiI-CM, Day 3: DiD) for histological reconstruction of the injection sites. Following each injection, the craniotomy was sealed with KwikCast, and the mice were allowed to recover for 30 mins in their home cage before behavioral testing. For single day injection paradigm, mice underwent head-fixation and microinjections as described with only the muscimol injection (100 nl Muscimol (5 mM) with CTB (1:1 v/v)) and no recovery session. In the testing phase, mice were presented with 50 stimulus trials and 50 catch trials and performance was quantified across the entire session on each day. After the final day of the testing phase, animals were anaesthetized and perfused, and the brain dissected for histological confirmation of injection sites (Shamash et al. 2018).

#### Acoustic perception task

The acoustic perception task followed the same procedure as thermal training, but the thermal stimulus was replaced by a 4 kHz, 85 dB, 2 s acoustic stimulus delivered from a speaker 10 cm from the mouse.

### Sensory stimulation

Thermal stimuli were applied to the plantar surface of the mouse forepaw. The paw was fixed to the center of the thermal stimulator using tape to ensure consistent contact with the active surface. Thermal stimuli of varying amplitude (22, 26, 28, 30, 31, 25, 28, 40, 41, 42°C) were delivered from an adapted temperature of 32°C using gold-plated Peltier elements (4 x 10 mm; 75°C/s; QST Lab). Thermal stimuli were delivered with an interstimulus interval of 18s. For behavioral experiments, thermal stimuli were delivered by a ceramic Peltier element (8 mm x 8 mm; 20°C/s) that was controlled by a custom feedback-controlled thermal stimulator (ESYS GmbH, Berlin). Thermal stimuli (27°C for cool training and 42°C for warm training were delivered every 20-30 s.

Vibrotactile stimuli (40 Hz, 2s) were provided using a force-feedback movement sensor arm (Aurora Scientific, Dual-Mode Lever Arm Systems 300-C) that was held in contact with the back of the forepaw. During vibrotactile stimulation, the paw was maintained at 32°C.

### Data analysis and statistics

#### Data pre-processing pipeline

Data pre-processing was performed in accordance with the Allen Institute Ecephys Spike Sorting Pipeline (https://github.com/AllenInstitute/ecephys_spike_sorting). Briefly, the spike-band data underwent four pre-processing steps before spike sorting: offset removal, median subtraction, filtering, and whitening. Kilosort2 (https://github.com/MouseLand/Kilosort) was utilized to identify spike times and assign them to individual units. Units that were not considered noise were organized into Neo-package files (https://pypi.org/project/neo/). A range of metrics were calculated to facilitate the identification of well-isolated single units that were included in the dataset and multiunit activity. These metrics are broadly separated into waveform-based parameters, which assess the physical properties of the units’ waveforms, and spiking-based parameters, which assess the firing properties and the isolation of each unit relative to other units from the same recording (as shown in Extended Data Fig. 1).

#### Spike analyses

Assessing thermo-tactile responsivity: For each unit, spike trains were binned in 100-ms bins, and the maximum spike-count bin within a 2-second baseline time window before stimulus onset was compared with spike-count bins within a 2-second time window during stimulus presentation using a Wilcoxon signed-rank test. A unit was classified as ‘responsive’ if the p-value of the statistical test was smaller than 0.013, respectively.

Population PSTH: Individual spike trains from units classified as responsive were binned in 5 ms bins and a population spiking matrix was created which was averaged across units and smoothed with a Gaussian Kernel (25 ms bandwidth). The resulting population spiking vectors were aligned to stimulus onset.

Temperature Tuning Index: The Temperature Tuning Index was calculated using the trial-averaged and baseline-corrected peak firing rate during 2 s stimulus duration at maximum cold and warm temperature amplitude:

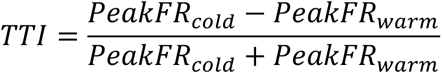

Units were classified into three categories: Cool, Cool-warm, and Warm using the TTI. TTI values between -0.3 and +0.3 were considered cool-warm while TTI values outside of this range were considered selective for cool or warm.

Response Latency: For each responsive unit, spike trains were binned in 1 ms bins and smoothed with a Gaussian Kernel with 10 ms bandwidth for cold stimuli and 100 ms for warm stimuli. The latency of the response defined as the peak of the 2nd derivative of the smoothed spike train between stimulus onset and peak of the 1st derivative.

Peak Response Time: For each responsive unit, spike trains were binned in 1 ms bins and smoothed with a Gaussian Kernel with 10 ms bandwidth for cold stimuli and 100 ms for warm stimuli. The peak response time was defined as the peak smoothed spike train.

Duration Index: The Temperature Duration Index (TDI) was calculated using the trial-averaged and baseline-corrected peak firing rate during the initial 1 s and final 1 s of stimulus presentation:

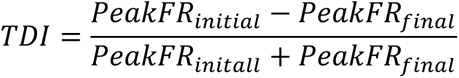

### Reconstruction of probe and unit sites

#### Probe tracks

The digital images were processed using the Matlab-based software, SharpTrack (https://github.com/cortex-lab/allenCCF). The individual images were first downsampled, cropped, and aligned to the Allen Institute Common Coordinate Framework (CCFv3). Probe points were then visually identified and marked. To account for variations in brain size and changes due to fixation, the precise positions of the probes were scaled based on several criteria. These included: 1) the relative low-frequency LFP power across channels, as a dramatic decrease in LFP power indicates the brain surface as well as decrease at the Hippocampus-Thalamus border 2) anatomical landmarks, such as ventricles or the corpus callosum, where low unit extraction was expected; and 3) the waveforms duration of units assigned to inhibitory structures in the thalamus, where the majority of cells have a short duration waveform. The resulting 3-dimensional coordinate of each electrode pad was entered as indices into the CCF reference volume to retrieve its estimated location in the mouse brain, which allowed estimation of single-unit locations. Visualization of probe trajectories was made using BrainRender (https://github.com/brainglobe/brainrender).

#### Spatial maps of responsive cells

Each single unit was assigned to the electrode contact at which the maximum waveform amplitude was registered, which allowed the reconstruction of the coordinate values in the anterior-posterior and dorsal-ventral axes of all recorded single units. A null distribution of all recorded cells in a particular region was used to normalize the cooling- or warming-responsive spatial distributions, respectively, to estimate the probability of a responsive cell in a defined spatial bin. If the number of neurons was insufficient to calculate the proportion of cooling or warming-responsive cells, the spatial bin was excluded from further analysis. Spatial maps showing the contour lines drawn at the 10^th^, 20^th^, 30^th^, and 40^th^ percentiles of the responsive population were overlayed with contours of thalamic structure in order to visualize the spatial distribution in a particular thalamic structure.

### Statistics

Sample sizes were not predetermined using statistical methods, but our sample sizes were similar to those used in previous publications. Trial order were pseudo-randomized. Descriptive statistics of distributions are presented as medians and interquartile ranges (IQR) unless otherwise noted. The statistical significance of differences between independent samples was evaluated using the non-parametric Kruskal-Wallis H test. Similarly, the significance of differences between dependent samples was assessed with the non-parametric Wilcoxon Rank Sum Test.

## Funding

European Research Council ERC-2015-CoG-682422 (JFAP)

Deutsche Forschungsgemeinschaft FOR 2143 (JFAP)

Deutsche Forschungsgemeinschaft SFB 1315 (JFAP)

National Institutes of Health NIH R01 NS123711 (JFAP)

Helmholtz Association (JFAP)

Human Frontiers Science Program LT000359/2018 (CW)

## Data and materials availability

All data and code are available upon request. We thank Sylvain Crochet and Jens Kremkow for advice and reading the manuscript, and Svenja Steinfelder and Nadine Groß for administrative and technical support.

## Extended Data

**Extended Data Figure 1.**
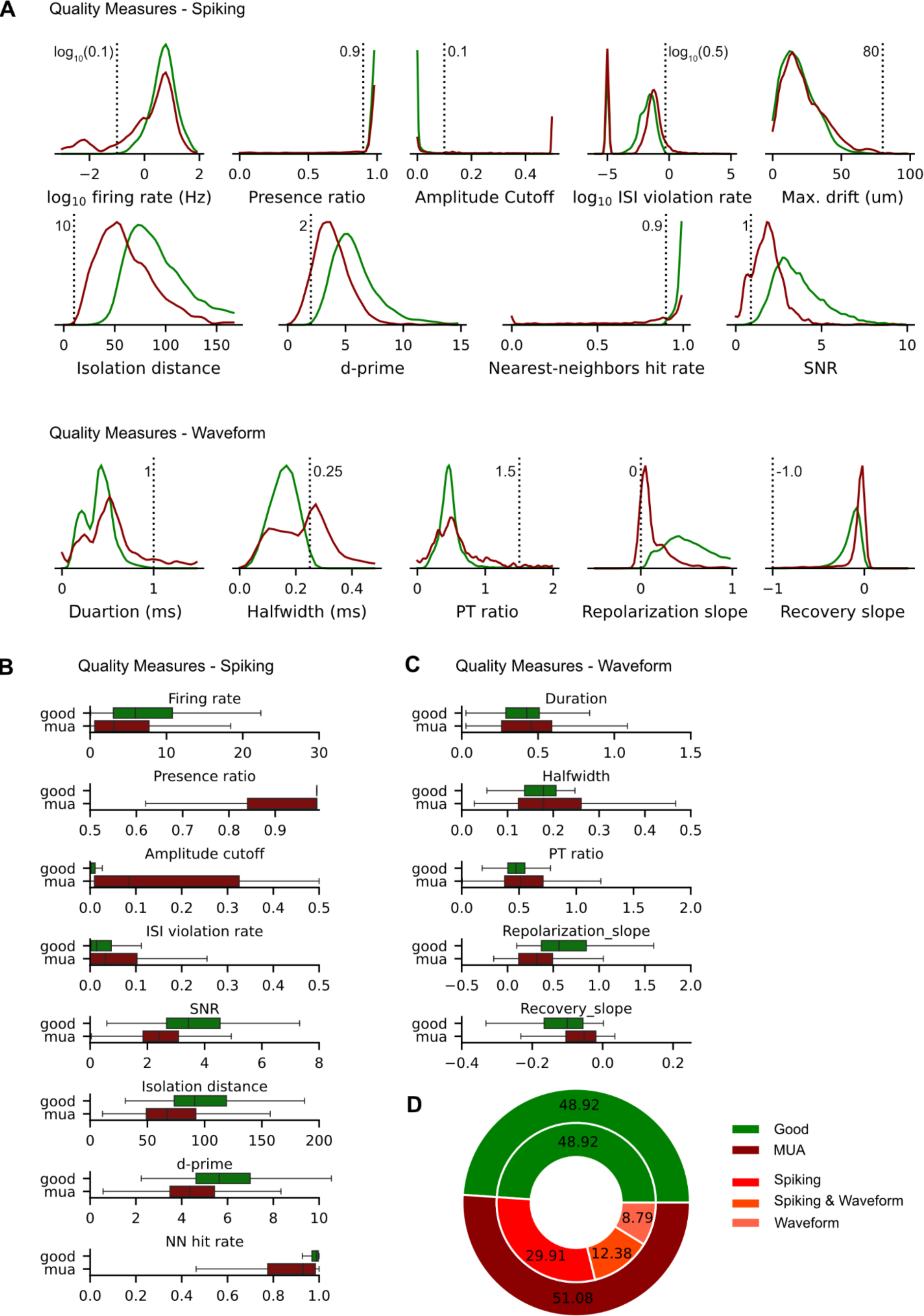
Spike quality metrics used to classify units. A. Spike quality metric distributions for all units classified as good (green) or as multi-unit activity (MUA; dark red). The threshold for good units is shown for each measure as a vertical dashed line. From left-to-right, top-to-bottom, the metrics shown are the spiking quality measures: unit firing rate, the presence ratio, the inter-spike-interval violation rate, the signal-to-noise ratio, the isolation distance, the d-prime value, the nearest neighbor hit rate; and the waveform quality measures: the waveform duration, the waveform half-width, the waveform peak-to-trough ratio, the repolarization slope, and the recovery slope. B. Box-and-whisker plots are shown for each of the spiking quality measures for good units (green) and MUA (dark red). C. Same as B, for waveform quality measures. D. Ratio of all recovered units classified as good units or MUA. The reason for classification as MUA is further subdivided into failure of passing the threshold on spiking quality measures, waveform quality measures, or both.

**Extended Data Figure 2.**
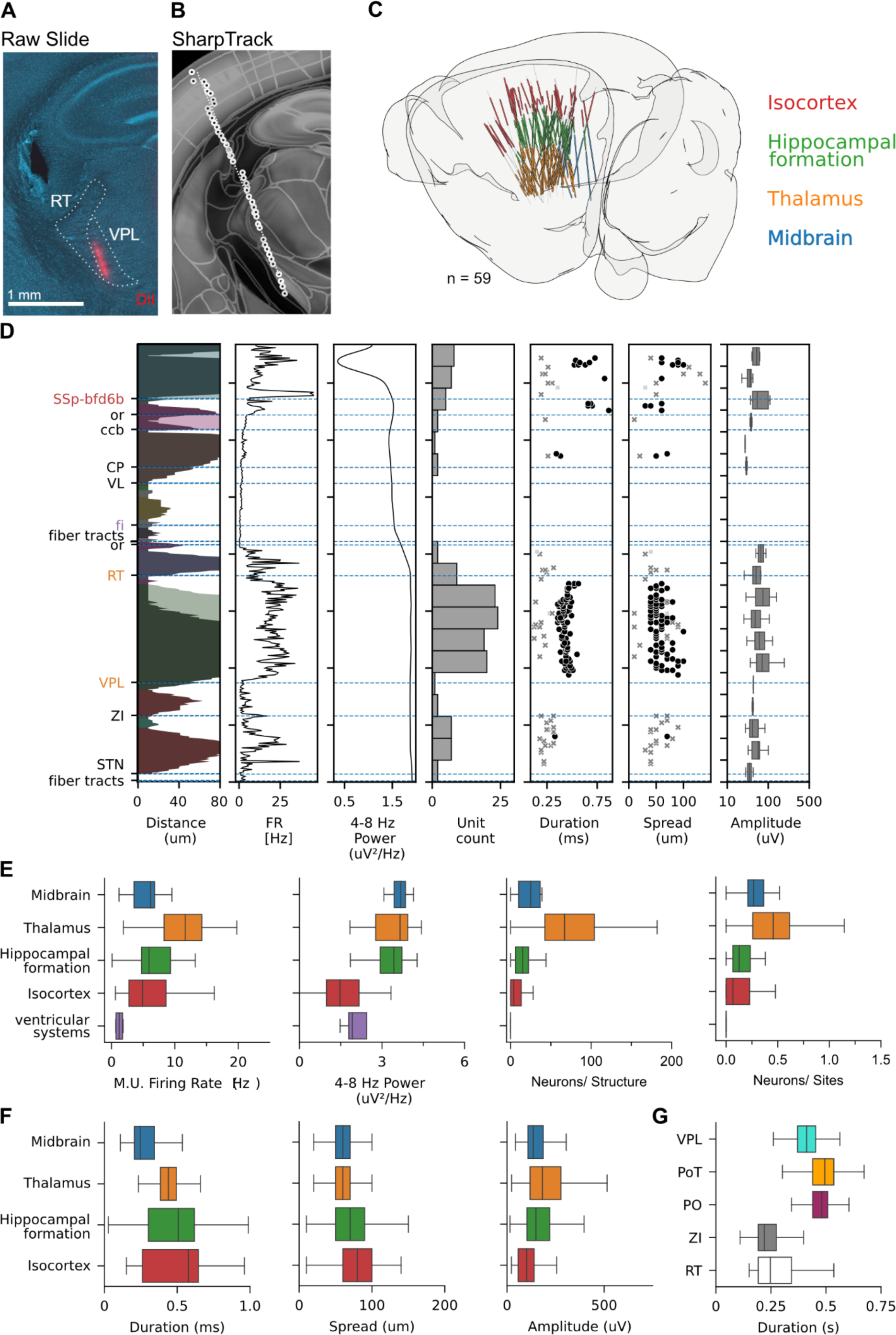
Anatomical reconstruction of Neuropixels probe trajectories. A. Example coronal brain slice with probe trajectory traveling through VPL (blue: DAPI, red: DiI). Scale bar: 1 mm. B. SharpTrack reconstruction of example mouse shown in panel A. Dots indicate reconstructed locations and dashed line depicts probe trajectory. C. Reconstructed probe trajectories for all recordings (n = 59 probes). Probe contacts are color-coded by gross anatomical structure. D. Example probe reconstruction parameterization depicting, from left-to-right, the estimated brain region, the firing rate, the low frequency power, the number of units, and three parameters of the waveform of the recovered units: the waveform duration, the spread across the probe, and the amplitude of the waveform. Data is shown for example in A/B. E. Distribution of probe reconstruction parameters within each major brain division across recordings (n = 59 recordings) from left to right: multiunit firing rate, low frequency power, neurons per structure, and neurons per site. F. Distribution of probe reconstruction parameters related to spiking waveforms within each major brain division across recordings (n = 59 recordings) from left to right: spike waveform peak-to-trough duration, spike waveform spread, and amplitude of the spike waveform. G. Spike waveform peak-to-trough ratio for thalamic nuclei that contain predominantly excitatory projection neurons (VPL, PoT, PO) and nuclei that provide thalamic inhibition (ZI, RT).

**Extended Data Figure 3.**
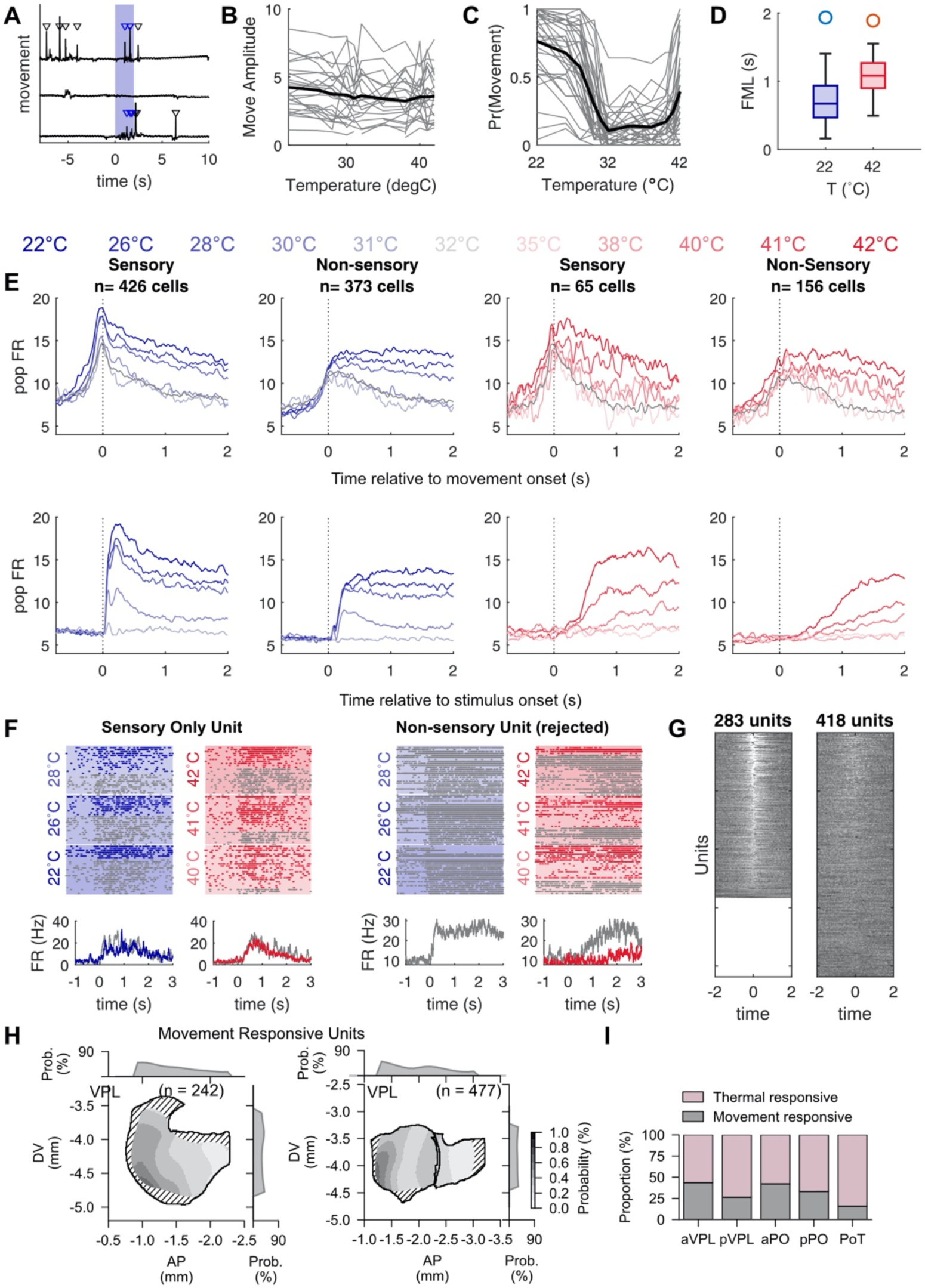
Thermal stimuli can evoke paw movements that drive thalamic activity in a minority of thermosensitive neurons. A. Example movement recording traces for three stimulus presentation trials (black). Thermal stimulus presentation (22°C) is indicated by the shaded blue bar. Detected movement events are indicated by the inverted triangle (blue triangle: movement during stimulus presentation; black triangle: movement outside of stimulus window). B. Amplitude of the movement events across stimulus intensities. Each gray line indicates a single session (n = 39 sessions, black line indicates mean). No relationship between movement intensity and stimulus amplitude. C. Probability of stimulus-evoked movement across stimulus intensities is strongly modulated by stimulus intensity (n = 39 sessions). D. First movement latency in response to cooling (22°C; median 0.674 seconds) and warming (42°C; median 1.110 seconds) stimulation (median +/- IQR). E. Population averaged peristimulus time histogram for cooling (blue) and warming (red) stimulation for units classified as sensory or non-sensory (see methods). Population PSTH on stimulus trials is aligned to movement onset (top) or stimulus onset (bottom). Note the graded response amplitude prior to movement onset for sensory units relative to non-sensory units, indicating they are not purely movement driven responses. F. Example raster and PSTH split by movement (grey) and non-movement trials (color). Left: Cooling stimulus evoked responses (grey: movement trials, blue: non-movement trials). Right: Warming stimulus evoked responses (grey: movement trials, red: non-movement trials) shown for a unit that has both a thermosensory response (left) and an excluded unit that has only a movement response (right). G. Peristimulus time histogram for each thermally responsive unit in response to a spontaneous movement event for units that are (left) and are not (right) driven by movement events. H. Spatial density maps for thalamic units responsive to movement in VPL (n = 242 movement responsive units) and in the PO complex (n = 477 movement responsive units). I. Proportion of movement responsive neurons by thalamic sub-region.

**Extended Data Figure 4.**
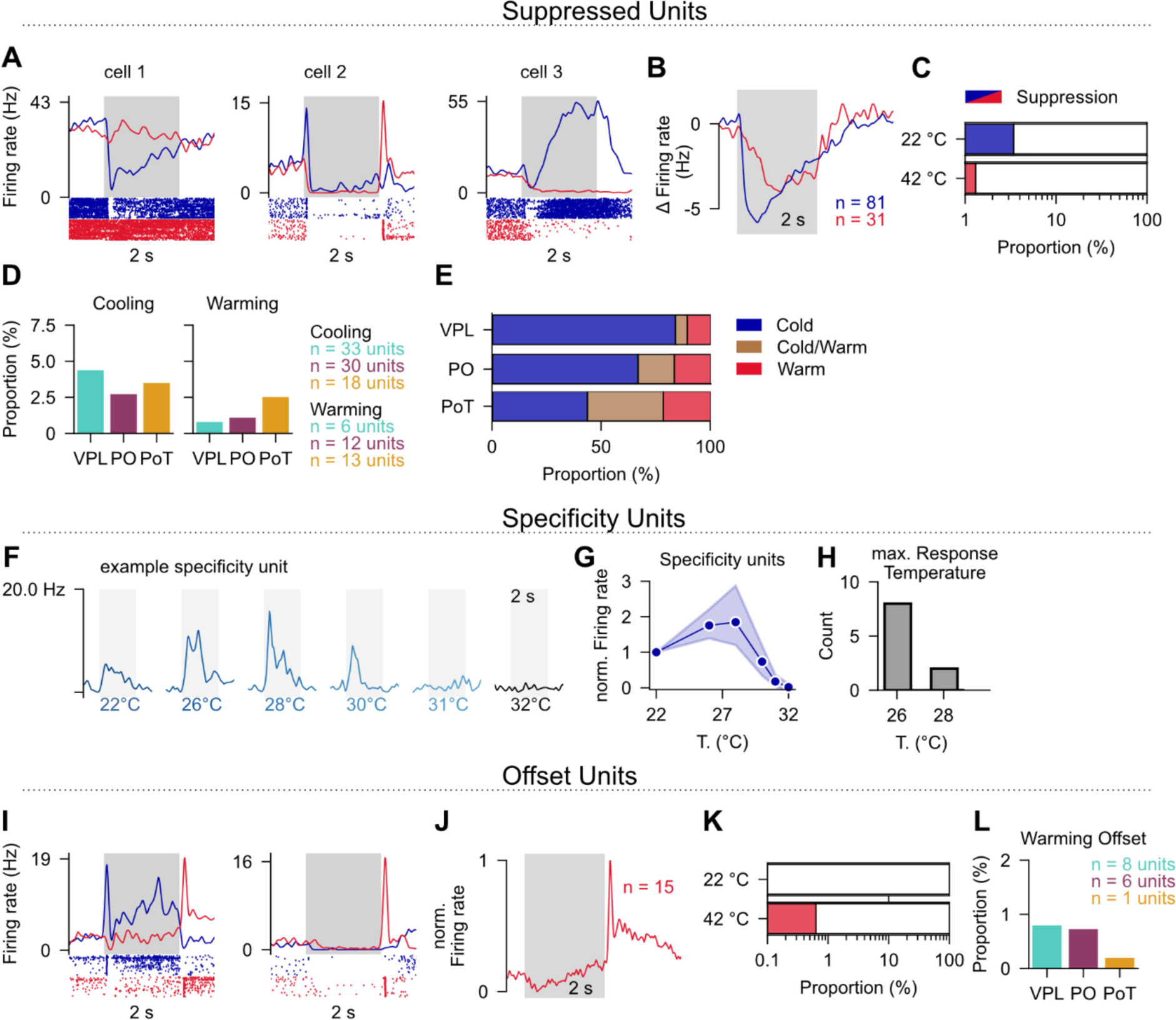
Characterization of minority thermosensory-evoked response types. A. Example thalamic neurons that show a suppressed response to thermal stimulation. B. Normalized stimulus-evoked PSTH to 22°C (blue, n = 81 neurons) and 42°C (red, n = 31 neurons) for all units with a suppressed stimulus evoked response. Stimulus duration indicated by grey shading. C. Proportion of responsive cells that are suppressed by cooling, or warming. D. Proportion of responsive cells that are suppressed by cooling (left) or warming (right) subdivided by thalamic nucleus. E. Proportion of suppressed responsive units classified Cool only, Cool/Warm, or Warm only response type by thalamic nucleus. F. Example cool-selective thermal specificity unit. G. Normalized firing rate for the example specificity units (n = 10 units), Filled circles show median, shaded region shows 95% confidence interval H. Number of units with a maximum response amplitude at 26°C or 28°C. I. Example thalamic neurons that show an offset thermosensory-evoked response. J. Normalized stimulus-evoked PSTH to 42°C (red) for all units with a stimulus-offset response. Stimulus duration indicated by grey shading. K. Proportion of responsive cells that are show an offset response. L. Proportion of responsive cells that are show a warming offset response subdivided by thalamic nucleus.

**Extended Data Figure 5.**
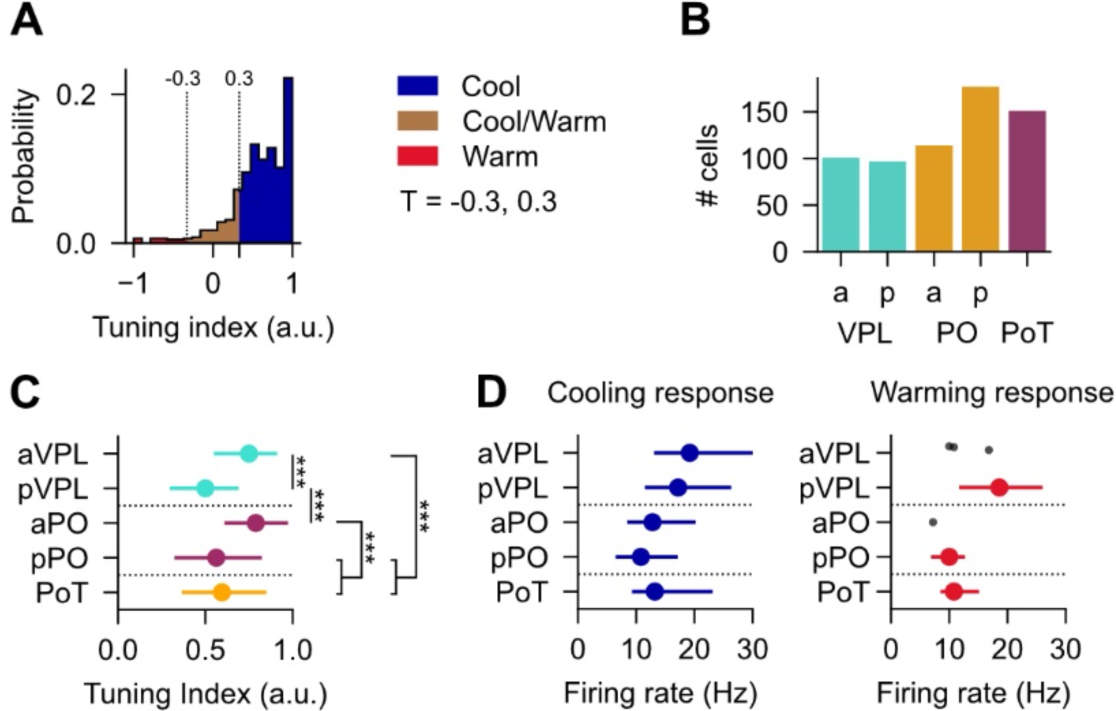
Cooling and warming representation across thalamic sub-nuclei. A. Thermal tuning index across responsive neurons. Cool, Warm or Cool/Warm response types are defined by the thermal tuning index (blue, copper, red, respectively). A tuning index value of [-1,-0.33] = Warm, [-0.33,0.33] = Cool/Warm, (0.33,1] = Cool. B. Number of cells located within each thalamic sub-nucleus classification. C. Average thermal tuning index (as computed in A) by thalamic sub-nucleus. Filled circles show median, error bars show IQR. Kruskal-Wallis H-test (*: p<0.05, **: p<0.01, ***: p<0.001) D. Cooling (left) and warming (right) evoked firing rate for each thalamic sub-nucleus. Filled circles show median, error bars show IQR

**Extended Data Figure 6.**
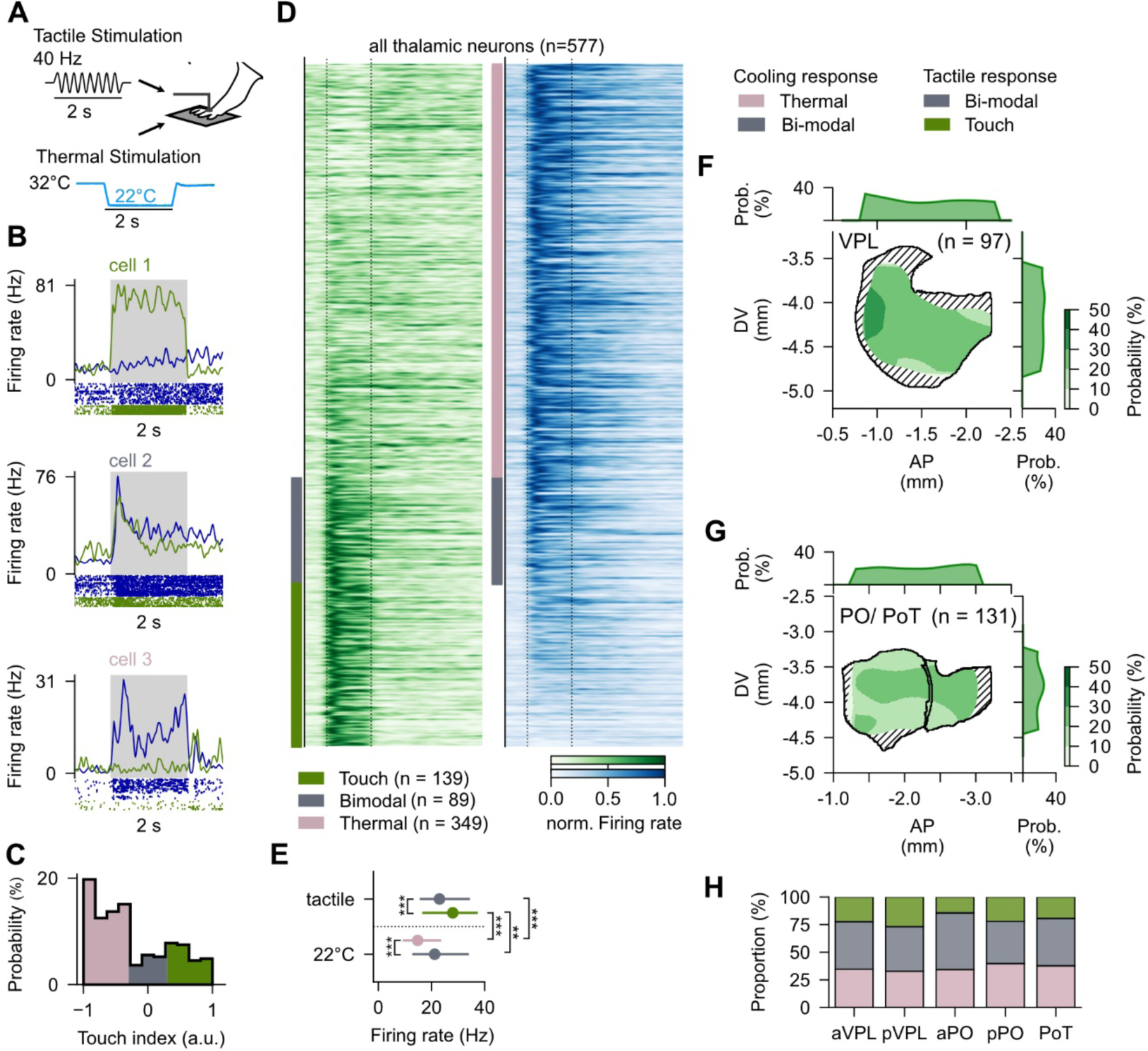
Comparable spatial location of touch and cool responsive neurons. A. Experimental schematic B. Example thermal (blue; cool) and tactile (green) sensory evoked responses (3 example units) C. Thermotactile tuning index across responsive neurons (n = 577 units). Classification as ‘thermal’, ‘mixed’, or ‘touch’ response type was defined by the thermotactile tuning index (pink, grey, green, respectively). A tuning index value of [-1,-0.33) = thermal, [-0.33,0.33] = mixed, (0.33,1] = tactile. D. Peristimulus time histogram for each thermotactile responsive unit in response to a tactile (green; left) or 22°C (right; blue) stimulus. Index bar indicates statistical significance for each unit as indicated in C (green: tactile n = 139 units, grey: mixed n = 89 units, pink: thermal n = 349 units) E. Tactile (left) and cooling (right) evoked firing rates for each thermotactile response type. Filled circles show median, error bars show IQR. Kruskal-Wallis H-test (*: p<0.05, **: p<0.01, ***: p<0.001) F. Spatial density maps for units responsive to touch in VPL (n = 97 touch units). G. Spatial density maps for units responsive to touch in the PO complex (n = 131 touch units). H. Proportion of thermotactile neurons by thalamic sub-region.

**Extended Data Figure 7.**
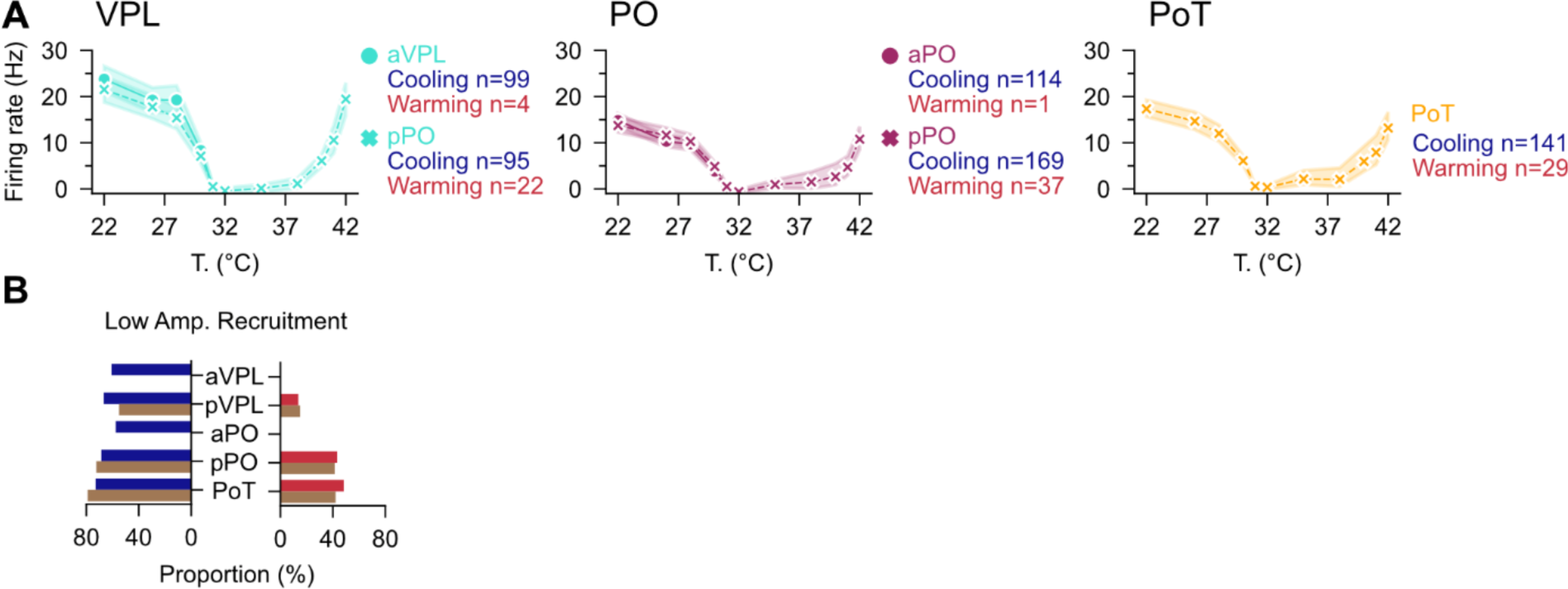
Further investigation of the thermal tuning parameters across thalamic nuclei and response types. A. Thermal tuning curves for each thalamic nucleus with VPL and PO subdivided into the anterior and posterior regions (aVPL: n = 99 cooling neurons, n = 4 warming neurons; pVPL: n = 95 cooling neurons, n = 22 warming neurons; aPO: n = 114 cooling neurons, n = 1 warming neurons; pPO: n = 169 cooling neurons, n = 37 warming neurons; PoT: n = 141 cooling neurons, n = 29 warming neurons). Filled circles show median, shaded show 95% confidence interval. B. Fraction of neurons that are recruited at low amplitudes shown for each region, subdivided by response type (cooling stimulus, left; warming stimulus, right).

**Extended Data Figure 8.**
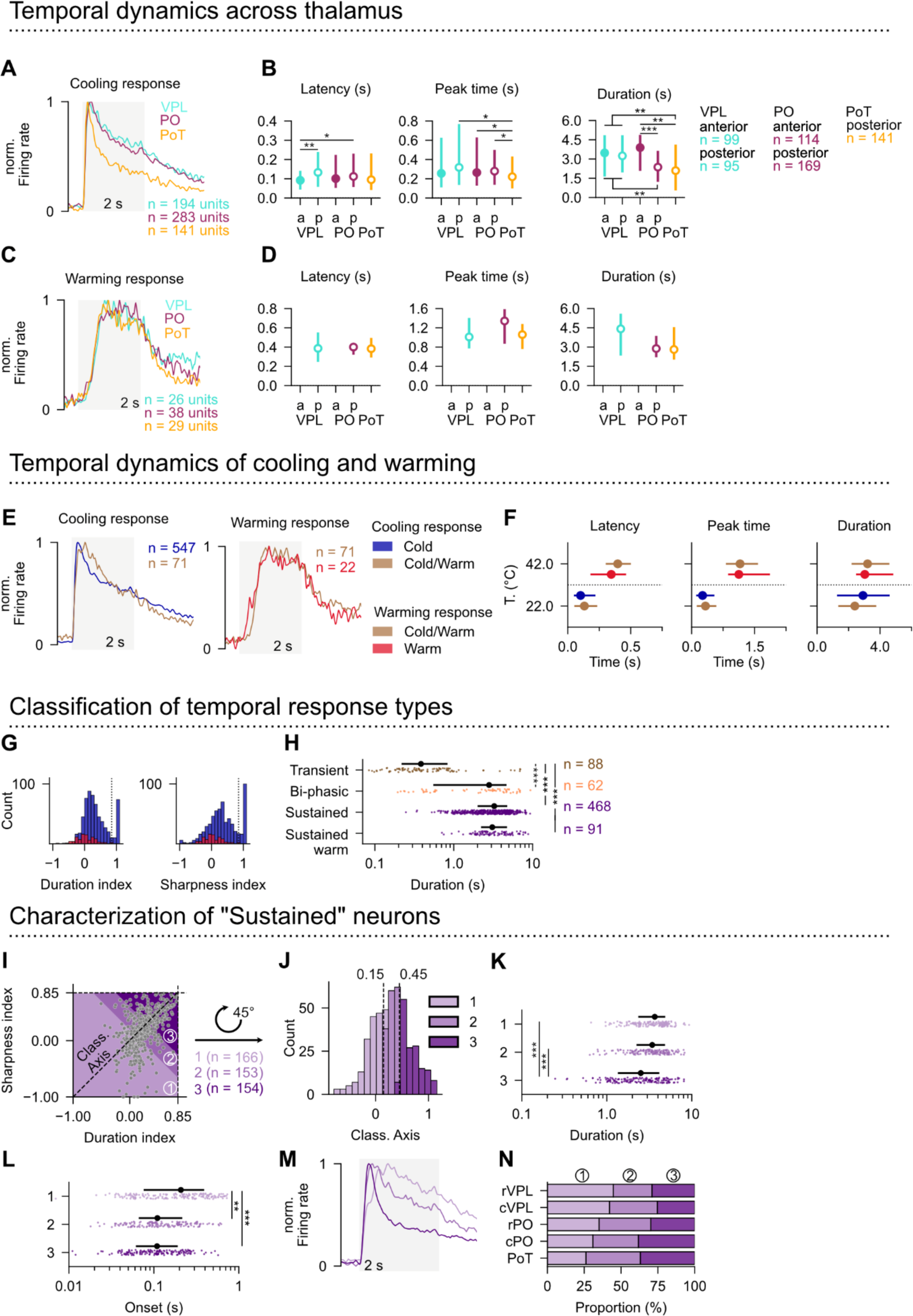
Further analysis of the dynamics of cooling and warming responses across the thalamus. A. Normalized stimulus-evoked PSTH by thalamic nucleus (top; VPL: teal, n = 194 neurons; PO: maroon, n = 283 neurons; PoT: gold, n = 141 neurons) to 22°C stimulation (bottom; stimulus trace). B. Stimulus response onset (left), peak time (middle), and duration index (right) in response to cool (22°C) by thalamic sub-nucleus. Statistical comparisons made using Kurskal-Wallis H-test (*: p<0.05, **: p<0.01, ***: p<0.001) Circles show median, error bars show IQR. C. Normalized stimulus-evoked PSTH by thalamic nucleus (top; VPL: teal, n = 26 neurons; PO: maroon, n = 38 neurons; PoT: gold, n = 29 neurons) to 42°C stimulation D. Stimulus response onset (left), peak time (middle), and duration index (right) in response to warming (42°C) by thalamic nucleus. Statistical comparisons made using Kurskal-Wallis H-test (*: p<0.05, **: p<0.01, ***: p<0.001) Circles show median, error bars show IQR. E. Normalized stimulus-evoked PSTH to 22°C (‘cold’ response type: n = 547 neurons, blue; ‘cold/warm’ response type: n = 71 neurons, copper) and 42°C (‘warm’ response type: n = 22 neurons, red; ‘cold/warm’ response type: n = 71 neurons, copper). Stimulus duration indicated by grey shading. F. Parameterized distributions for stimulus response onset (left), peak time (middle), and duration index (right) in response to cool (22°C) and warm (42°C, red) for ‘cold’, ‘cold-warm’, and ‘warm’ response types. Circles show median, error bars show IQR. G. Distribution of duration index (left) and sharpness index (right) for all cooling (blue) and warming (warm) responses across all thalamic neurons. H. Scatter plots display the duration as a function of response dynamic type. Dots denote individual units (transient: brown, n = 88 units; mixed: orange, n = 62 units; sustained: purple, n = 468 units, warming sustained: purple, n = 91) Black dots show median, error bars show IQR. I. Segmentation of the sustained response type into three subsections (1: n = 166 neurons, 2: n = 153 neurons, 3: n = 154 neurons). J. Cell position along classification axis determines subsection classification. K. Scatter plots display the duration for each sustained response dynamic quadrant in response to a cooling stimulus (22°C). Dots denote individual units (1: n = 166 neurons, 2: n = 153 neurons, 3: n = 154 neurons) and black dots denote the population distribution (median +-IQR). Statistical comparisons made using Kurskal-Wallis H-test (*: p<0.05, **: p<0.01, ***: p<0.001). L. Scatter plots display the onset latency for each sustained response dynamic quadrant in response to a cooling stimulus (22°C). Dots denote individual units (1: n = 166 neurons, 2: n = 153 neurons, 3: n = 154 neurons) and black dots denote the population distribution (median ± IQR). Statistical comparisons made using Kurskal-Wallis H-test (*: p<0.05, **: p<0.01, ***: p<0.001) M. Normalized cooling stimulus-evoked PSTH for each sustained response dynamic subsection (1: n = 166 neurons, 2: n = 153 neurons, 3: n = 154 neurons). Stimulus duration indicated by grey shading. N. Proportion of cooling sustained responsive units classified as subsection 1-3 by thalamic nucleus.

**Extended Data Figure 9.**
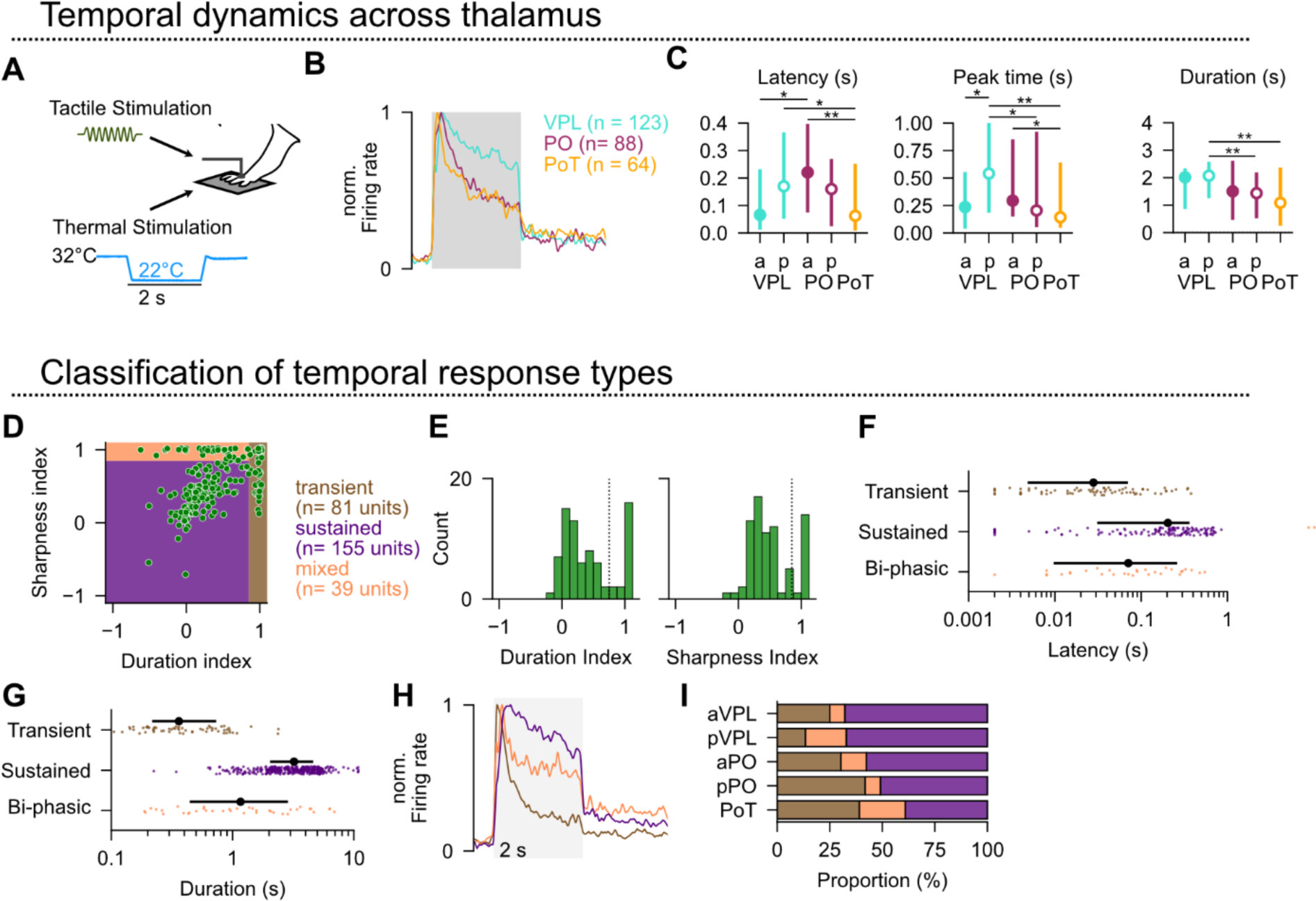
Tactile responses are more transient in posterior regions. A. Experimental schematic. B. Normalized tactile stimulus-evoked PSTH by thalamic nucleus (top; VPL: teal, n = 123 neurons; PO: maroon, n = 88 neurons; PoT: gold, n = 64 neurons). C. Stimulus response onset (left), peak time (middle), and duration index (right) in response to tactile stimulation by thalamic nucleus (top; VPL: teal, n = 123 neurons; PO: maroon, n = 88 neurons; PoT: gold, n = 64 neurons). Statistical comparisons for plots are shown only if significant. Filled circles show median, error bars show IQR. D. Scatterplot of the touch-evoked response dynamics parameterized as the duration index and the sharpness index (n = 275 neurons). Transient responses have a duration index > 0.85; Mixed response types have a sharpness index > 0.85 (and duration index < 0.85). Sustained responses have a duration & sharpness index < 0.85. E. Distribution of duration index (left) and sharpness index (right) for tactile responsive neurons (n = 275 neurons) F. Scatter plots display the latency for each response dynamic type in response to a tactile stimulus. Dots denote individual units (transient: brown, n = 84 neurons, mixed: orange, n = 43 neurons, sustained: purple, n = 148 neurons) and black dots denote the population distribution (median ± IQR). G. Scatter plots display the duration index for each response dynamic type in response to a tactile stimulus. Dots denote individual units (transient: brown, n = 84 neurons, mixed: orange, n = 43 neurons, sustained: purple, n = 148 neurons) and black dots denote the population distribution (median ± IQR). H. Normalized stimulus-evoked PSTH to touch stimulation for each response dynamic type. (transient: brown, n = 81 neurons, mixed: orange, n = 39 neurons, sustained: purple, n = 155 neurons). Stimulus duration indicated by grey shading. I. Proportion of tactile responsive units classified as displaying transient (brown), mixed (orange), or sustained (purple) response dynamics by thalamic sub-nucleus.

**Extended Data Figure 10.**
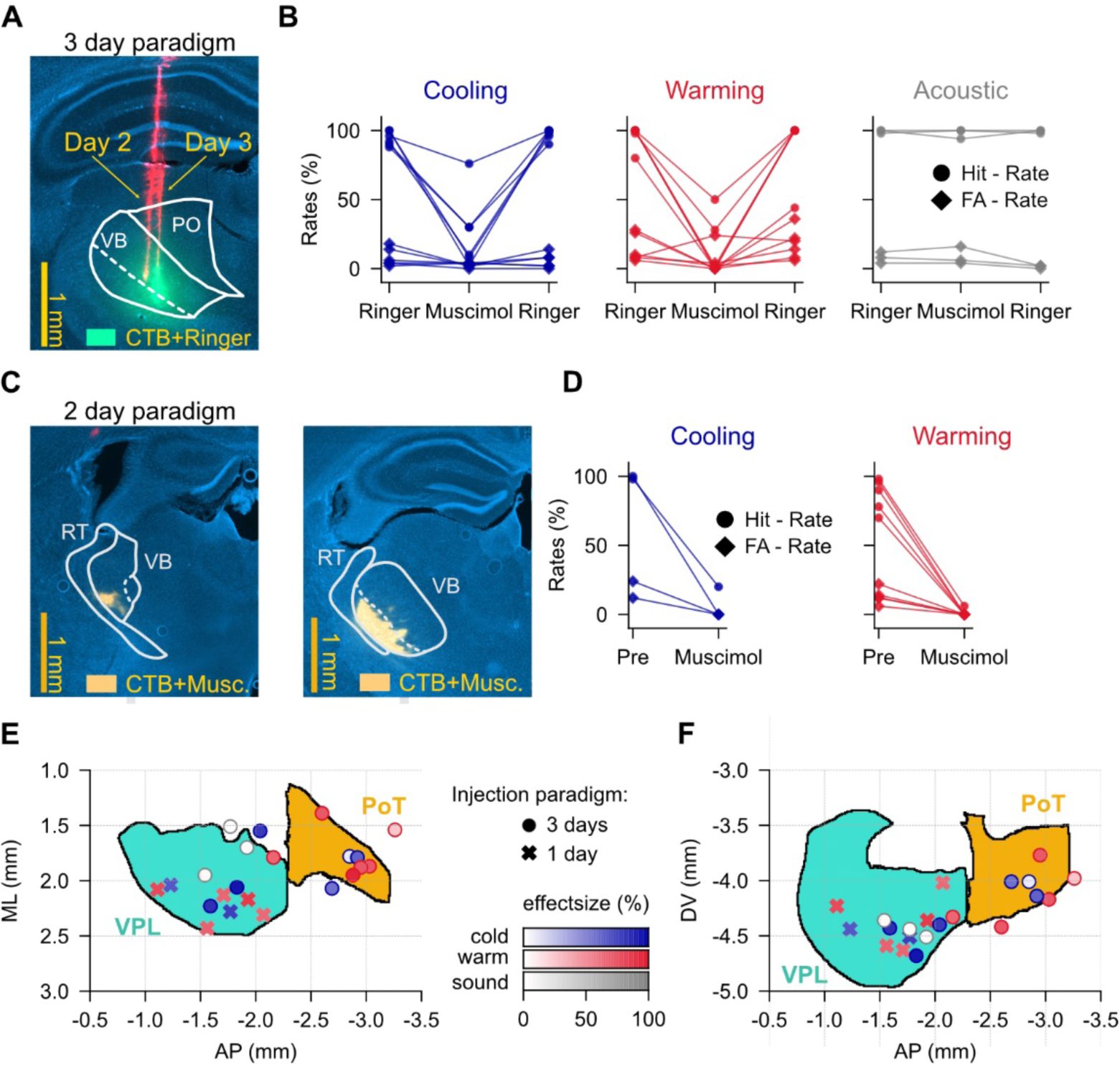
Extended data for thalamic inactivation during behavior. A. Example histological image depicting the DiI stained injection pipette tracts from Day 2 & 3 (red) and CTB (green) that was injected with ringer on Day 3 in DAPI (blue) labeled coronal section. Scale bar: 1 mm. B. Behavioral performance across three-day injection paradigm for mice trained on cooling (left, blue; n = 6 mice), warming (center, red; n = 6 mice), or auditory (left, grey: n = 3 mice) tasks shown as the hit rate (circle) and false alarm rate (diamond). C. Example histological image depicting the CTB (yellow) that was injected with muscimol on Day 2 in DAPI (blue) labeled coronal section. Scale bar: 1 mm. D. Behavioral performance across one-day injection paradigm for mice trained on cooling (left, blue; n = 2 mice) or warming (center, red; n = 5 mice) tasks shown as the hit rate (circle) and false alarm rate (diamond). E. Reconstructed position of the pipette tip in horizontal plane. Color indicates the trained task (cool: blue, warm: red), opacity indicates effect size (100% opaque: 100% effect; 0% opaque: 0% effect) and marker-shape indicates experimental paradigm (circle: three-day injection paradigm, cross: one-day injection paradigm). F. Same as E, but for the sagittal plane.

